# Pre-marked chromatin and transcription factor co-binding shape the pioneering activity of Foxa2

**DOI:** 10.1101/607721

**Authors:** Filippo M. Cernilogar, Stefan Hasenöder, Zeyang Wang, Katharina Scheibner, Ingo Burtscher, Michael Sterr, Pawel Smialowski, Sophia Groh, Ida M. Evenroed, Gregor D. Gilfillan, Heiko Lickert, Gunnar Schotta

**Author notes:** Corresponding authors: Gunnar Schotta, Heiko Lickert, Filippo Cernilogar. These authors contributed equally to this work (Zeyang Wang, Katharina Scheibner).

## Abstract

Pioneer transcription factors (PTF) can recognize their binding sites on nucleosomal DNA and trigger chromatin opening for recruitment of other non-pioneer transcription factors. However, critical properties of PTFs are still poorly understood, such as how these transcription factors selectively recognize cell type-specific binding sites and under which conditions can they can initiate chromatin remodelling. Here we show that early endoderm binding sites of the paradigm PTF Foxa2 are epigenetically primed by low levels of active chromatin modifications in embryonic stem cells (ESC). Priming of these binding sites is supported by preferential recruitment of Foxa2 to endoderm binding sites compared to lineage-inappropriate binding sites, when ectopically expressed in ESCs. We further show that binding of Foxa2 is required for chromatin opening during endoderm differentiation. However, increased chromatin accessibility was only detected on binding sites which are synergistically bound with other endoderm transcription factors. Thus, our data suggest that binding site selection of PTFs is directed by the chromatin environment and that chromatin opening requires collaboration of PTFs with additional transcription factors.

## Introduction

Transcription factors (TFs) drive lineage-specific transcription programs by binding gene regulatory elements dispersed throughout the genome (Spitz and Furlong, 2012). However, since DNA is wrapped around histones to form nucleosomes and chromatin, TFs have to overcome this physical barrier to bind their DNA target sites (Jiang and Pugh, 2009; Voss and Hager, 2014). Although most TFs can recognize their target sequence only on nucleosome-free DNA, so-called pioneer transcription factors (PTFs) have the peculiar ability to engage their target sequence on nucleosomal DNA (Iwafuchi-Doi and Zaret, 2014; Soufi et al., 2015). Following binding to their target sites, PTFs can induce chromatin opening supporting the recruitment of non-pioneer TFs and ultimately leading to activation of the underlying gene regulatory elements (Cirillo et al., 2002; Zaret et al., 2016). Interestingly, despite their potentially universal targeting, PTFs only bind to a subset of their potential DNA binding motif containing target sites (Donaghey et al., 2018; Hurtado et al., 2011; Zaret et al., 2016). These findings imply that additional mechanisms, such as cell-type specific cofactors (Liu and Kraus, 2017; Swinstead et al., 2016a) and chromatin environment (Lupien et al., 2008; Petruk et al., 2017; Soufi et al., 2012; Wang et al., 2015) can influence binding site selection of PTFs.

While it is widely recognized that PTFs have the capacity to engage with previously inaccessible regions of chromatin, there is still scarce understanding of how they initiate remodelling and opening of the surrounding chromatin. Binding of PTFs can lead to eviction of nucleosomes (Li et al., 2012a) or displacement of linker histone H1 (Iwafuchi-Doi et al., 2016). However, it is currently unclear how PTFs assemble distinct chromatin remodelling machineries on specific binding sites.

We have tackled those questions by studying the paradigm PTF Foxa2 in the physiological context of *in vitro* endoderm differentiation from mouse ESCs. We found that Foxa2 binding during endoderm differentiation is dynamic with stable and differentiation stage-specific binding sites. Endoderm-specific Foxa2 binding sites feature low levels of active chromatin modifications in ESCs, suggesting an epigenetic priming for Foxa2 recruitment during differentiation. We found that Foxa2 binding is required but not sufficient for chromatin opening. Rather, co-binding of Foxa2 with additional endoderm TFs appears necessary for chromatin opening. In summary, our data suggest that binding sites for pioneer transcription factors are epigenetically primed and that chromatin opening requires synergistic binding of transcription factors in close vicinity.

## Results

### Isolation of Foxa2 expressing mesendoderm and endoderm cells

To study Foxa2 binding site selection and Foxa2-dependent chromatin changes we decided to investigate the transition from pluripotent ESCs via mesendoderm (MESEND) progenitors to definitive endoderm (DE) cells using an *in vitro* differentiation system (Figure 1A). For isolating cells from specific stages of endoderm commitment in high purity we made use of a double knock-in (DKI) mouse ESC line carrying Foxa2-Venus and Sox17-Cherry fusion reporter genes (Burtscher et al., 2013; Burtscher et al., 2012). The resulting TF fluorescent fusion proteins are expected to be fully functional, as homozygous *Foxa2*^*FVF/FVF*^; *Sox17*^*SCF/SCF*^ mice are fully viable without any obvious phenotypes. By fluorescence-activated cell sorting (FACS) we isolated pure populations of Foxa2-Venus^neg^/Sox17-Cherry^neg^ pluripotent ESCs, Foxa2-Venus^pos^/Sox17-Cherry^neg^ mesendoderm cells after three days and Foxa2-Venus^pos^/Sox17-Cherry^pos^ cells after five days of differentiation (Fig 1A, Fig S1A). We refer to these cell populations as d0 (pluripotent ESCs), d3F (Foxa2^pos^ mesendoderm cells) and d5FS (Foxa2^pos^/Sox17^pos^ definitive endoderm cells). Transcriptome analyses of these populations revealed stage-specific expression signatures with 1053 differently expressed genes (fold change > 2; padj < 0.05; Fig 1B, Fig S1B,C, Table S1). Consistent with the progressive differentiation to endoderm we observed downregulation of pluripotency genes, a transient expression of mesoderm genes in d3F cells and progressive induction of endoderm genes in d3F and d5FS cells (Fig 1C). Expression of anterior endoderm markers (Cer1, Dkk1) and absence of posterior definitive endoderm markers (Cdx2) suggests that the *in vitro* differentiation favors the generation of cells resembling anterior definitive endoderm. To highlight the transcription factor network responsible for endoderm differentiation we identified the most influential transcription factors by their expression change and connectivity (Fig 1D). This analysis suggests that transcription factors such as Foxa2, Gata4 and Eomes appear as most important to initiate endoderm differentiation (d0-d3F network), while the importance of additional TFs emerges in later stages of endoderm differentiation (d0-d5FS network).

**Figure 1.**
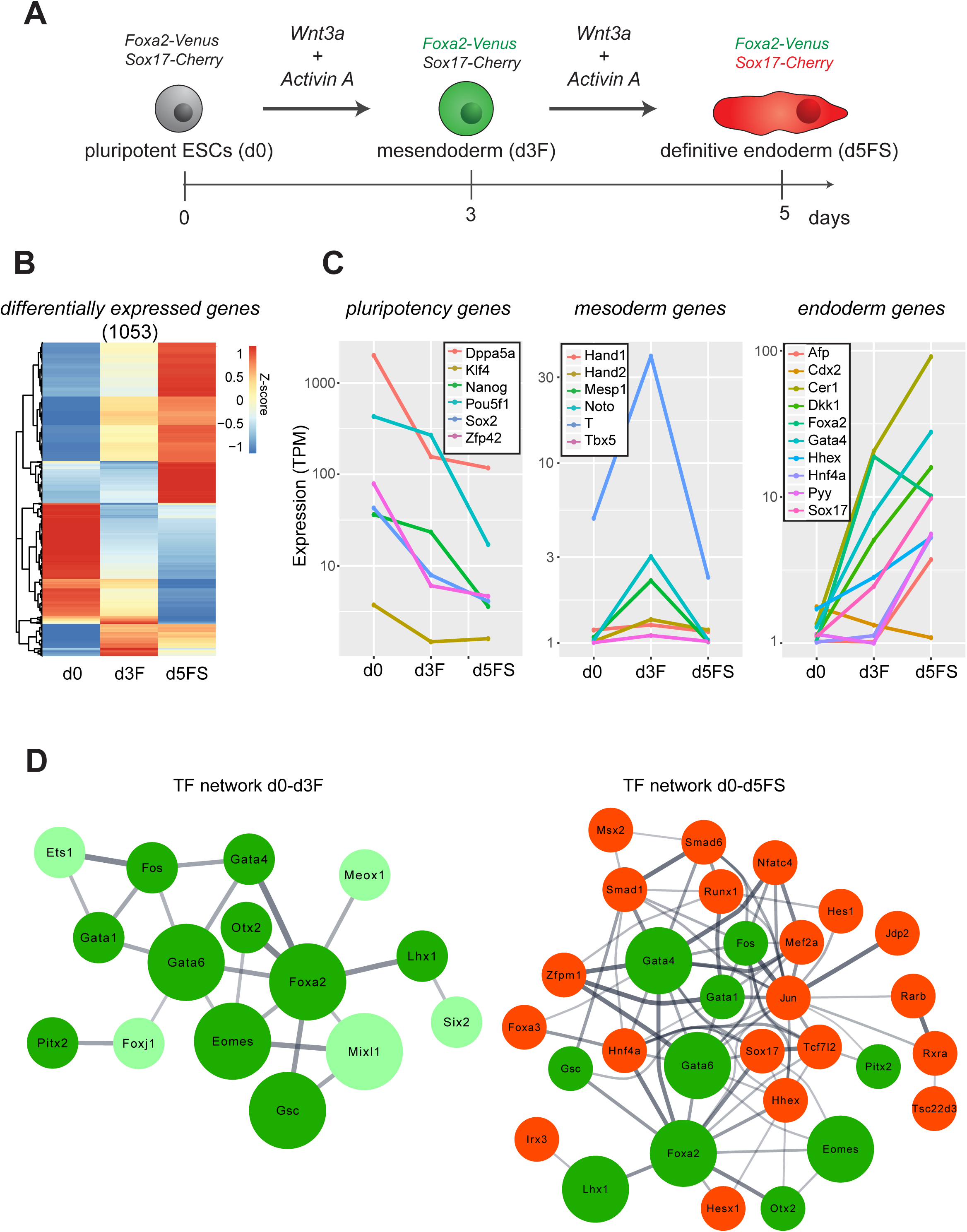
An in vitro differentiation system modelling early endoderm differentiation. A) Endoderm differentiation of ESCs is triggered by Wnt3A/Activin A treatment. Mesendoderm (Foxa2-Venus^+^; Sox17-Cherry^-^) and endoderm (Foxa2-Venus^+^; Sox17-Cherry^+^) cells can be isolated by FACS. B) Heat map showing z-scores of the expression levels of the 1053 differentially expressed genes between pluripotent ESCs (d0), mesendoderm (d3F) and endoderm (d5FS) cells (padj < 0.05, fold change > 2; n=2 for each condition). C) Average expression levels of selected marker genes for pluripotent, mesoderm and endoderm cells in the *in vitro* differentiated ESCs (TPM - Transcripts Per Kilobase Million; n=2 for each condition). D) Network of most influential transcription factors, driving transition from pluripotent ESCs (d0) through a mesendoderm stage (d3F) to definitive endoderm (d5FS) cells. Bigger nodes correspond to the top 5 transcription factors. Width of edges corresponds to String database (StringDB) scores. Only connected nodes are plotted. Color code: light green factors are specific to the d0-d3F network, green factors are present in both networks, red factors emerge in the d0-d5FS network.

Thus, by combining *in vitro* endoderm differentiation with FACS sorting we could isolate two consecutive stages of endoderm differentiation resembling features of mesendoderm and anterior definitive endoderm cells.

### Foxa motifs are over-represented in regions of increased chromatin accessibility upon endoderm differentiation

To get additional insight into the gene regulatory network governing the transition from mouse pluripotent ESCs via MESEND progenitors to the DE stage we used the assay of transposase-accessible chromatin using sequencing (ATAC-seq) (Buenrostro et al., 2013) to determine the genome-wide chromatin accessibility landscape in d0, d3F and d5FS cells. Overall, we identified 190606 accessible regions, located primarily at non-promoter regions (Fig S2A). Remarkably, the PCA analysis of all ATAC peaks, promoter peaks or non-promoter peaks demonstrated that in particular non-promoter ATAC peaks are strongly distinguishing features of the three cell populations (Fig S2C). We then assessed differential accessibility between the differentiation stages (fold change > 2, padj < 0.05) and found that 5.8% ATAC peaks change in the transition d0-d3F and 23.4% in the transition d0-d5FS (Fig 2A-C). The differentially accessible regions (DARs) are located primarily at non-promoter regions (Fig 2A). A heatmap of the top regulated ATAC peaks during endoderm differentiation (Fig 2B) recapitulates the pattern of transcriptional changes observed by RNA-seq (Fig 1B). Thus, DARs show a good correlation with regulated genes (Fig S2B), although the analysis is limited by connecting individual ATAC peaks with specific genes only by proximity to the TSS. To enhance the biological insights obtained from DARs we analysed the annotations of the nearby genes with the GREAT tool (McLean et al., 2010). In particular, peaks that change in the d0-d3F transition are associated with gene ontology annotations connected to loss of stem cell properties and the emergence of differentiated cells (Fig S2D-F, Table S2).

**Figure 2.**
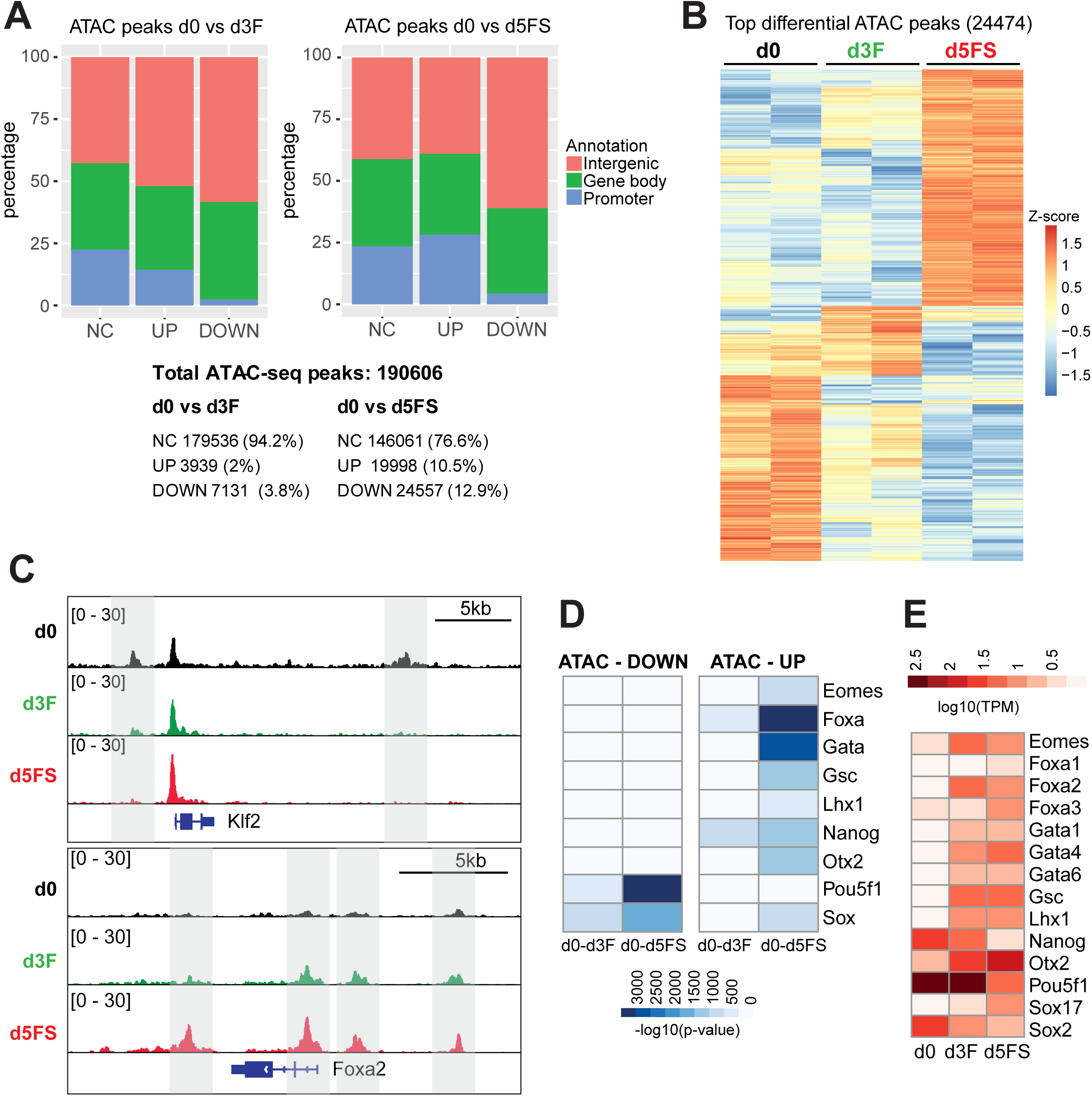
Foxa motifs are over-represented in regions of increased chromatin accessibility upon endoderm differentiation. A) Numbers and percentages of not changed (NC) and dynamic (UP/DOWN) ATAC-seq peaks associated with different genomic features in d0 vs d3F (left panel) or in d0 vs d5FS (right panel) comparison. Peaks are considered dynamic with an ATAC-seq coverage fold change >2. B) Heat map showing relative chromatin accessibility (z-scores of normalized ATAC-seq signals) of the top dynamic ATAC-seq peaks (24474) in pluripotent ESCs (d0), mesendoderm (d3F) and endoderm (d5FS) cells (padj < 0.05, fold change > 4; n=2 for each condition). C) Representative genome browser view of ATAC-seq signals in d0, d3F and d5FS cells. D) Heat map showing the p-values of transcription factor motif enrichments in dynamic ATAC-seq peaks. The Homer tool was used to scan for known motifs of expressed transcription factors (TPM > 1). Only the top scoring motifs with -log10 (p-value) > 500 are shown. Since members of Foxa, Gata and Sox families bind very similar motifs only the family names are given. Columns represent analyses for differential ATAC peaks between d0-d3F and d0-d5FS. E) Expression heatmap of transcription factors shown in (D) in d0, d3F and d5FS cells. Relevant Foxa, Gata and Sox family members are shown.

Next, to identify transcription factors responsible for establishing DARs during endoderm differentiation we determined DNA binding motifs within differential ATAC peaks (Table S3). We found that DARs with reduced accessibility (DOWN) are mostly enriched for motifs of pluripotency related TFs, such as Oct4 and Sox family factors (Fig 2D,E), consistent with downregulation of the pluripotency network. In contrast, DARs with increased accessibility (UP) are mostly enriched with motifs of mesendoderm- and endoderm-related TFs, such as Foxa family, Gata family, Eomes, Lhx1, Otx2 and Gsc (Fig 2D,E). Notably, upregulated DARs most prominently feature Foxa motifs. Based on the expression of the Foxa family members (Fig 2E) and the fact that Foxa2 is expressed first among the Foxa1-3 family (Kaestner, 2010) we predict that the PTF Foxa2 is a key factor to induce chromatin accessibility in the context of endoderm differentiation.

### Loss of Foxa2 impairs endoderm differentiation

Foxa2 knock-out mice show early embryonic lethality (E9-E10) and absence of anterior definitive endoderm and axial mesoderm (Ang and Rossant, 1994; Weinstein et al., 1994), indicating its functional importance for mesendoderm and endoderm development. To investigate if Foxa2 is also critical for *in vitro* endoderm differentiation, we generated a Foxa2 knock-in/knock-out allele (Foxa2^Venus^) by replacing the coding region of Foxa2 with an H2B-Venus expression cassette in mouse ESCs (Fig S3A-C). As H2B-Venus is under control of the Foxa2 promoter we detected nuclear H2B-Venus protein only upon mesendoderm differentiation in both control (Foxa2^Venus/+^) and Foxa2 ko (Foxa2^Venus/Venus^) ESCs, suggesting that initiation of Foxa2 expression is independent of Foxa2 protein (Fig S3D).

Next, we investigated Foxa2-dependent transcriptional changes. We induced endoderm differentiation in both control and Foxa2^Venus^ ko ESCs and FACS-isolated Venus-positive cells at day 3 of differentiation (Fig 3A, Fig S3E). We then performed RNA-seq based transcriptome analysis of undifferentiated (control = d0^con^; ko = d0^ko^) and differentiating (control = d3^con^, ko = d3^ko^) cells. In agreement with the fact that Foxa2 is not expressed in ESCs we did not observe transcriptional differences between d0^con^ and d0^ko^ cells, however, differentiating d3^con^ and d3^ko^ cells were clearly distinct (Fig S3F). We identified 1268 differently expressed genes (fold change > 2, padj < 0.01) between d3^con^ and d3^ko^ cells (Fig 3B, Table S4). Gene ontology analysis of these genes showed a dominant enrichment for terms associated with embryonic development (Fig S3G). We found that in d3^ko^ cells pluripotency genes were not properly downregulated and endoderm genes were not fully activated, while mesoderm genes did not show obvious changes (Fig S3H-L). These findings reflect downregulation of an endoderm signature gene set (Grapin-Botton, 2008) in d3^ko^ cells (Fig 3D). The failure to differentiate to endoderm is likely to be linked with the aberrant transcription factor network of d3^ko^ cells (Fig 3C), which has no overlap with the endoderm differentiation networks observed in d3F and d5FS cells (Fig 1D).

**Figure 3.**
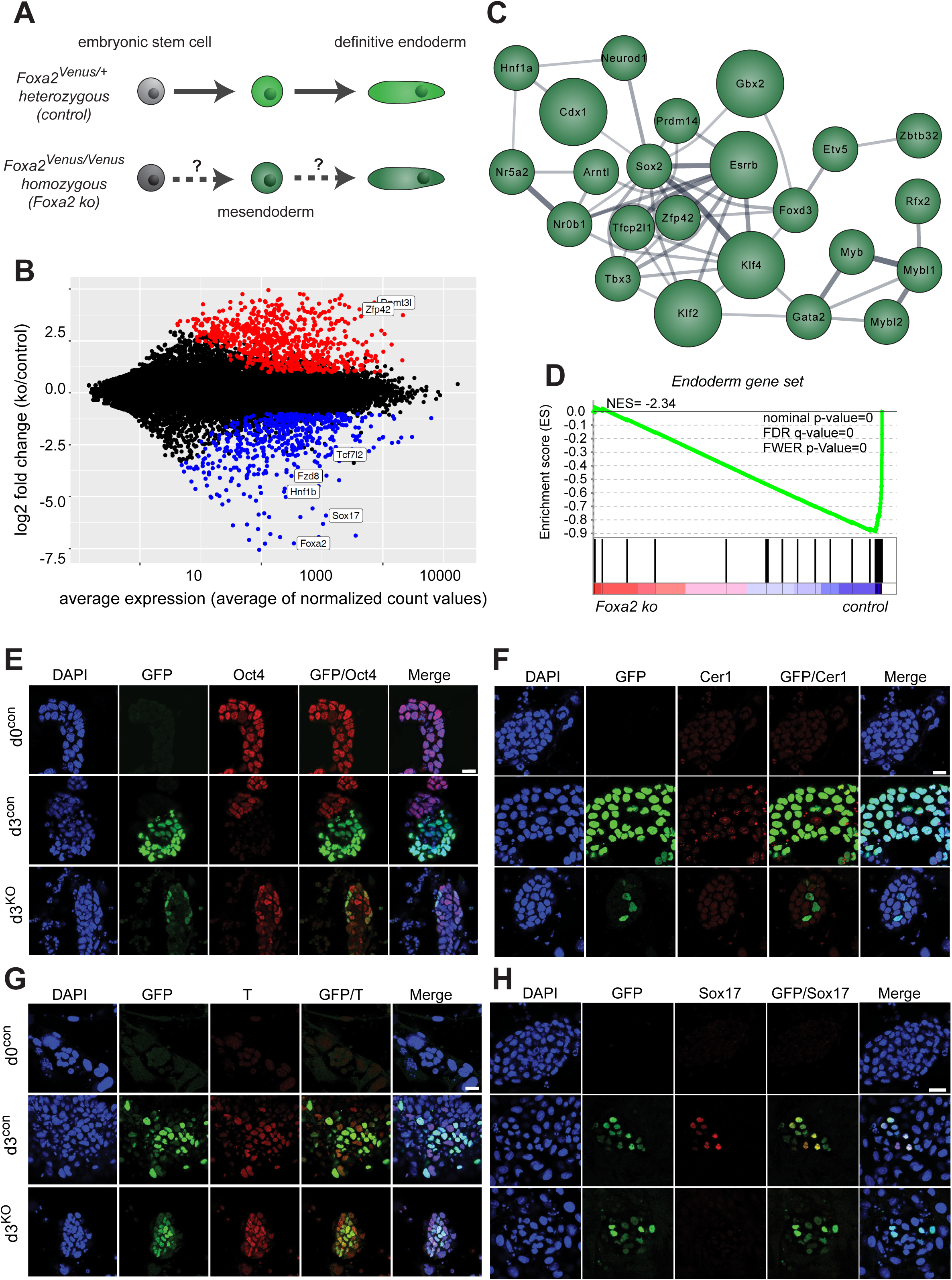
Loss of FOXA2 impairs endoderm differentiation. A) Differentiation and FACS sorting strategy of control vs. Foxa2 ko ESC into endoderm. B) Dotplot showing average expression vs. log2-fold change of coding genes in endoderm differentiating Foxa2^Venus/+^ control vs. Foxa2^Venus/Venus^ ko cells. Coloured dots indicate genes with significantly changed expression (adjusted p-value < 0.05, fold change > 2; n=3 for each condition). Positions of relevant genes are indicated. C) Network of most influential transcription factors in endoderm differentiating Foxa2^Venus/Venus^ ko cells. Bigger nodes correspond to the top 5 transcription factors. Width of edges corresponds to String database (StringDB) scores. The network has no overlap with the one shown in Fig. 1D. D) Gene set enrichment analysis (GSEA) of an endoderm gene set between control (d3^con^) and Foxa2 ko cells (d3^ko^). The Foxa2 ko cells show strong underrepresentation of these endoderm signature genes. NES: normalized enrichment score. E) Confocal sections showing undifferentiated Foxa2^Venus/+^ (d0^con^), endoderm differentiating Foxa2^Venus/+^ (d3^con^) and Foxa2^Venus/Venus^ homozygous (d3^KO^) cells stained with antibodies to Venus/GFP (green), Oct4 (red) and DAPI (blue). Scale bar: 20µm. F) Confocal sections showing undifferentiated Foxa2^Venus/+^ (d0^con^), endoderm differentiating Foxa2^Venus/+^ (d3^con^) and Foxa2^Venus/Venus^ (d3^KO^) cells stained with antibodies to Venus/GFP (green), Brachyury/T (red) and DAPI (blue). Scale bar: 20µm. G) Confocal sections showing undifferentiated Foxa2^Venus/+^ (d0^con^), endoderm differentiating Foxa2^Venus/+^ (d3^con^) and Foxa2^Venus/Venus^ (d3^KO^) cells stained with antibodies to Venus/GFP (green), Cer1 (red) and DAPI (blue). Scale bar: 20µm. H) Confocal sections showing undifferentiated Foxa2^Venus/+^ (d0^con^), endoderm differentiating Foxa2^Venus/+^ (d3^con^) and Foxa2^Venus/Venus^ (d3^KO^) cells stained with antibodies to Venus/GFP (green), Sox17 (red) and DAPI (blue). Scale bar: 20µm.

Consistent with the transcriptional changes we detected sustained levels of Oct4 protein d3^ko^ cells (Fig 3E). Mesendoderm marker gene Brachyury (T) was comparable between d3^con^ and d3^ko^ cells (Fig 3G), whereas key endoderm TFs and signaling factors, such as Cer1 and Sox17, were not induced in d3^ko^ cells (Fig 3F,H). In summary, our expression analyses show that Foxa2 ko cells are not able to fully activate the endoderm program which likely results in appropriate downregulation of important pluripotency genes. Taken together, these data demonstrate that Foxa2 is a master regulator for endoderm differentiation in ESC *in vitro* differentiation comparable to its in vivo function (Burtscher and Lickert, 2009).

### Foxa2 binding sites are dynamic during endoderm differentiation

To gain better insight into the fundamental roles of Foxa2 for endoderm differentiation we mapped Foxa2 binding sites by chromatin immunoprecipitation followed by next generation sequencing (ChIP-seq) in d3F and d5FS cells. We made use of a specific Foxa2 antibody (Tsankov et al., 2015) and considered only those sites common to two replicates as high confidence binding sites. Interestingly, Foxa2 binding is highly dynamic between the differentiation states. We identified 3411 binding sites specific for d3F cells, 4271 binding sites which are shared between d3F and d5FS and 3446 binding sites specific to d5FS cells (Fig 4A,D, Table S5). We named these categories of binding sites “transient”, “stable” and “late”, respectively. All binding categories display prominent presence of the Foxa DNA motif (Fig. S4A, Table S6), demonstrating specificity of the ChIP experiment. Consistent with previous reports (Alder et al., 2014; Wederell et al., 2008) we observed the majority of Foxa2 binding sites located at non-promoter regions (Fig 4B), suggesting that Foxa2 is primarily involved in gene regulation through distal cis-regulatory regions. Furthermore, to gain insights into the biological function of the Foxa2 binding sites, we analysed the annotations of the nearby genes with the GREAT tool (McLean et al., 2010). All the three categories of Foxa2 binding sites are enriched for gene ontology annotations associated to differentiation and development (Fig S4B). Stable and late Foxa2 binding sites are also flanked by genes of the Foxa network, but only stable binding sites are enriched for genes of the Wnt pathway (Fig S4C), suggesting a diversified biological function for the different Foxa2 binding sites. Next we aimed for correlating Foxa2 binding with gene expression changes. Due to the large number of Foxa2 binding sites not all these binding events are likely to cause changes in gene expression, as also observed for other transcription factors (Yamamizu et al., 2016). However, we found that a large percentage (∼30%) of genes that change expression over the time course or in Foxa2 ko cells are bound by Foxa2 (Fig S4D,E), suggesting that Foxa2 is important for regulating their expression and consistent with the important role of Foxa2 for endoderm development.

**Figure 4.**
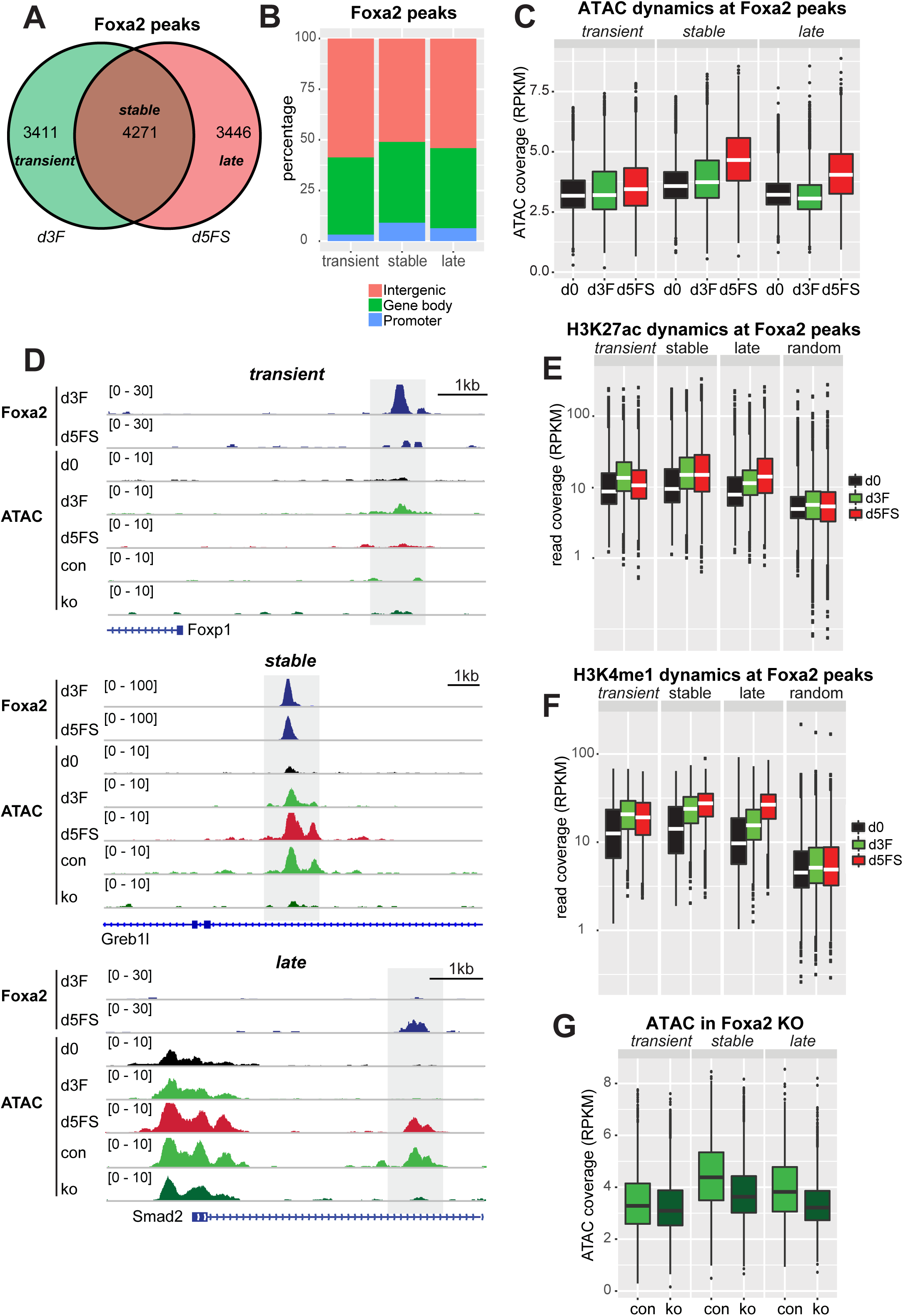
Foxa2 is required for chromatin opening. A) Venn diagram of Foxa2 binding sites in d3F (green) and d5FS (red) cells. d3F- and d5FS-specific binding sites were assigned “transient” and “late”, respectively; overlapping binding sites were assigned “stable”. n=2 for each condition. B) Percentages of transient, stable and late Foxa2 binding site associated with different genomic features. C) Box plot of normalized ATAC-seq coverage on transient, stable and late Foxa2 binding sites in d0, d3F and d5FS cells. n=2 for each condition. Wilcoxon ranks-sum test statistics is shown in Table S7. D) Genome browser view of examples for transient, stable and late Foxa2 binding sites. The following tracks are displayed: Foxa2 ChIP-seq in d3F and d5FS cells; ATAC-seq in d0, d3F, d5FS, Foxa2^Venus/+^ (con) and Foxa2^Venus/Venus^ (ko) endoderm differentiating cells. Dashed regions indicate Foxa2 binding sites. E) Box plot of normalized H3K27ac ChIP-seq coverage on Foxa2 binding sites and random genomic regions in d0, d3F and d5FS cells. n=2 for each condition. Wilcoxon ranks-sum test statistics is shown in Table S7. F) Box plot of normalized H3K4me1 ChIP-seq coverage on Foxa2 binding sites and random regions in d0, d3F and d5FS cells. n=2 for each condition. Wilcoxon ranks-sum test statistics is shown in Table S7. G) Box plots of normalized ATAC-seq coverage on transient, stable and late Foxa2 binding sites in endoderm differentiating Foxa2^Venus/+^ (con) and Foxa2^Venus/Venus^ (ko) cells isolated at day 3 of differentiation. n=3 for each condition. Wilcoxon ranks-sum test statistics is shown in Table S7.

Taken together, our data show that Foxa2 binding is highly dynamic during endoderm differentiation, with transient, stable and late binding sites in the vicinity of key developmental genes.

### Foxa2 is required for chromatin opening and recruitment of active histone modifications

Being a pioneer factor, Foxa2 is expected to mediate chromatin opening (Iwafuchi-Doi et al., 2016) or nucleosome depletion (Li et al., 2012a; Zaret et al., 2016) on its binding sites. We wondered if the different categories of Foxa2 binding sites show differences in these respects and functions. Thus, we analysed ATAC-seq coverage on transient, stable and late Foxa2 binding sites as a proxy for chromatin accessibility. Remarkably, we detected major differences in chromatin accessibility (Fig 4C,D, Table S7). In stable and late Foxa2 binding sites we could observe increased chromatin accessibility mainly in d5FS cells (Fig 4C,D, Fig S4G-I). In contrast, transient Foxa2 binding sites showed almost no change in chromatin accessibility (Fig 4C,D, Fig S4F,I). These data demonstrate that binding of Foxa2 in d3F cells does not lead to increased chromatin accessibility at most of its binding sites.

Since Foxa2 binding did not fully correlate with increased chromatin accessibility, we asked if Foxa2 recruitment would better correlate with changes in enhancer chromatin modifications (Creyghton et al., 2010). For both, H3K4me1 and H3K27ac, we observed increased levels on stable and late binding sites during endoderm differentiation (Fig 4E,F, Table S7). Enhancer modifications were also increased in d3F cells on transient binding sites, however, loss of Foxa2 from these binding sites in d5FS cells correlated with reduced levels of these modifications (Fig 4E,F, Fig S4L). These data demonstrate that Foxa2 binding strictly correlates with establishment of enhancer chromatin modifications, but only on a subset of binding sites, increased chromatin accessibility can be induced.

To understand if increased chromatin accessibility depends on Foxa2 binding we performed ATAC-seq in Foxa2 deficient cells. As Foxa2^Venus^ ko cells do not carry the *Sox17*^*SCF*^ allele we were not able to isolate specific populations corresponding to d3F and d5FS, but rather FACS-isolated endoderm differentiating cells based on Foxa2 expression. Comparing control (Foxa2^Venus/+^) with Foxa2 ko (Foxa2^Venus/Venus^) cells, we detected reduced chromatin accessibility on stable and late Foxa2 binding sites, whereas no substantial differences could be observed on transient binding sites (Fig 4D,G, Table S7, Fig S4F-I). In summary our data indicate that Foxa2 is required but not sufficient for chromatin opening at its binding sites.

### Co-binding of Foxa2 with other TFs correlates with chromatin opening

We found that binding of Foxa2 to a target locus is not sufficient to induce chromatin accessibility. Therefore, we hypothesized that co-binding of additional proteins, probably other transcription factors, may favour chromatin accessibility. To assess this hypothesis, we analysed the presence of endoderm TF binding motifs at Foxa2 binding sites. As expected, the Foxa motif is strongly enriched in transient, stable and late binding sites (Fig 5A, Table S6). Motifs of other mesendoderm- and endoderm-related TFs, such as Gata family, Lhx1 and Gsc, tend to be enriched on stable and late Foxa2 binding sites which display increased chromatin accessibility (Fig 5A). Transient binding sites at which Foxa2 fails to induce chromatin accessibility showed no enrichment for these additional TF binding sites. The presence of a binding motif is not necessarily predictive of actual binding. Thus, we generated ChIP-seq profiles for Gata4, a prominently expressed member of the Gata family, in d3F and d5FS cell populations. Consistent with the motif predictions our analysis revealed that Gata4 peaks coincide with Foxa2 peaks preferentially at stable and late Foxa2 binding sites (Fig 5B,C, Fig S5A). Most of these binding sites display strongly increased chromatin accessibility during endoderm differentiation (Fig S5B). Collectively, these data support a model wherein cooperative TF binding and activity is necessary to induce chromatin opening.

**Figure 5.**
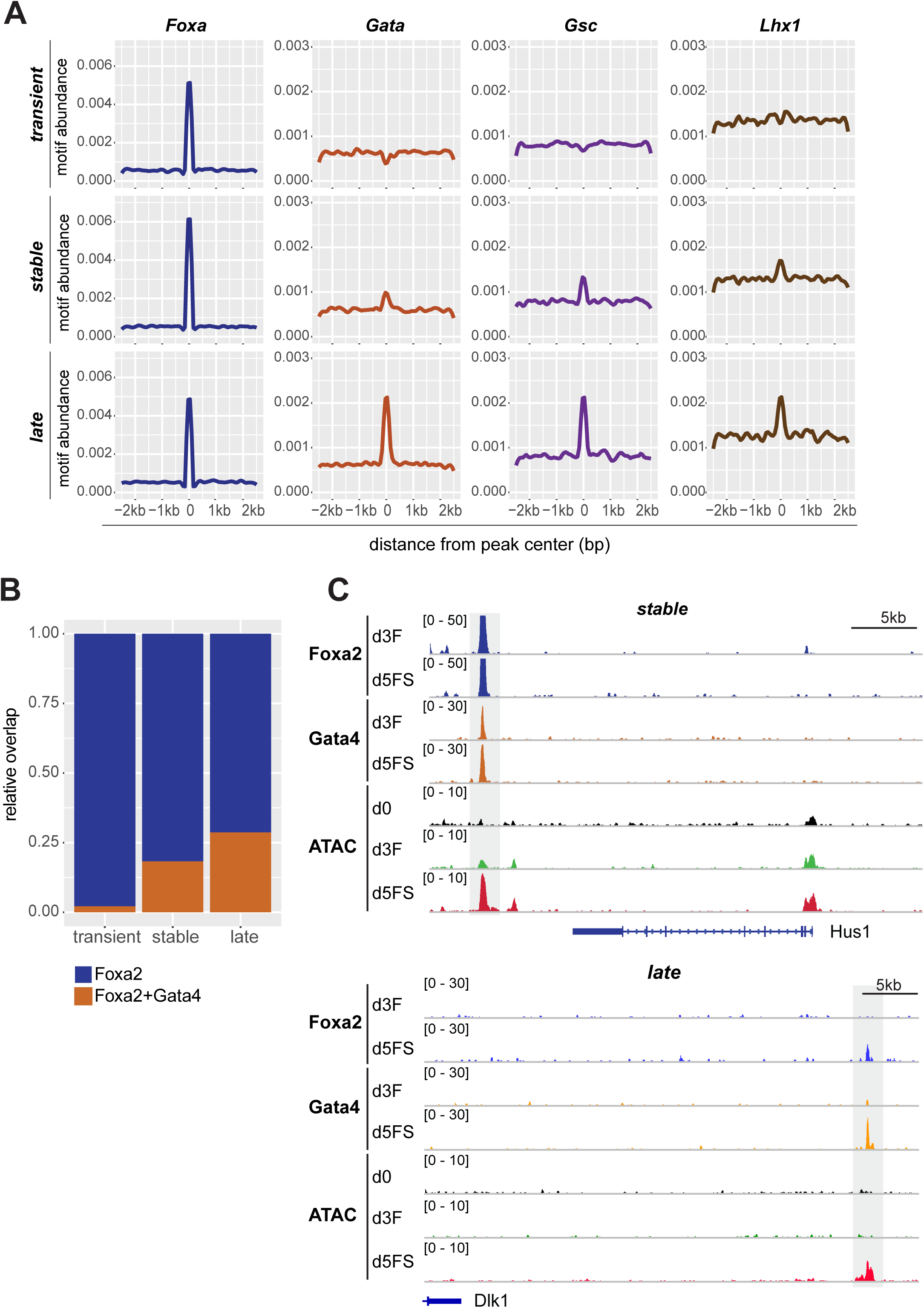
Co-binding of Foxa2 with other TFs correlates with chromatin opening. A) Density plots for motif abundances of Foxa, Gata, Gsc and Lhx1 motifs in transient, stable and late Foxa2 binding sites. B) Fraction of transient, stable and late Foxa2 binding sites bound by Foxa2 (blue) or co-bound by Foxa2 and Gata4 (brown). C) Genome browser view of example stable and late Foxa2 binding sites. The following tracks are displayed: Foxa2 and Gata4 ChIP-seq in d3F and d5FS cells, ATAC-seq in d0, d3F and d5FS cells. Dashed regions indicate stable (upper panel) and late (lower panel) Foxa2 binding sites.

### Endoderm-specific Foxa2 binding sites feature active chromatin modifications in ESCs

Foxa2 is continuously expressed during endoderm differentiation in endoderm, pancreatic and liver progenitors as well as in differentiated insulin-producing beta cells (Kaestner, 2010). However, Foxa2 only binds a subset of its potential binding sites and clear binding differences exist between cell types (Donaghey et al., 2018). As chromatin environment could influence transcription factor binding (Lupien et al., 2008; Petruk et al., 2017; Soufi et al., 2012), we thought to determine chromatin modifications in ESCs which might distinguish endodermal from other Foxa2 binding sites bound at later stages during differentiation. For this analysis we investigated endodermal Foxa2 binding sites (transient, stable and late) compared with pancreatic beta cell-specific binding sites from a published dataset (Jia et al., 2015). For comparison we defined Nanog binding sites representative of active regulatory regions, Trim28 binding sites (Castro-Diaz et al., 2014) corresponding to repressed chromatin, and random genomic regions (Fig 6A).

**Figure 6.**
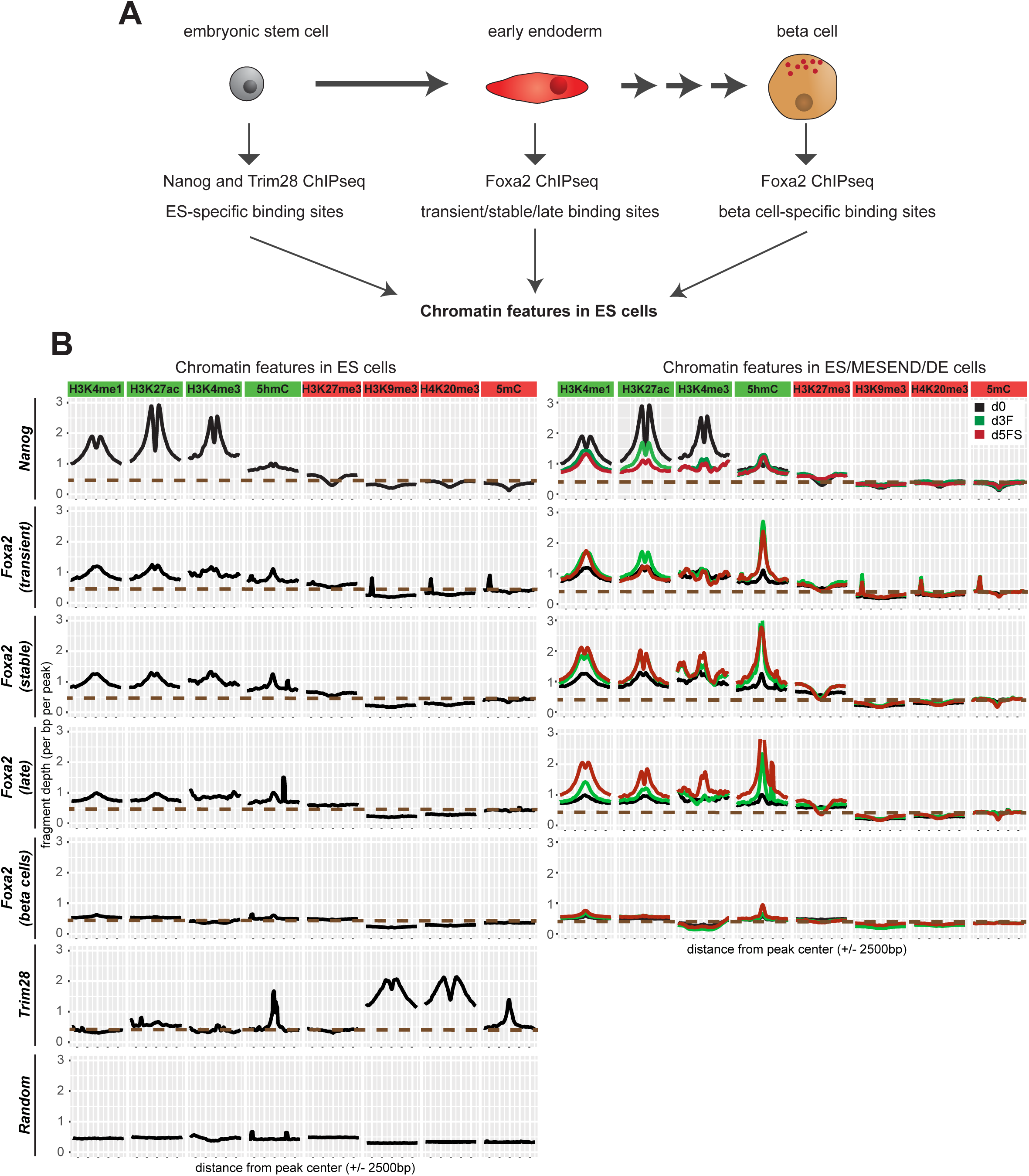
Endoderm-specific Foxa2 binding sites feature active chromatin modifications in ES cells. A) Embryonic stem cells can differentiate in definitive endoderm cells which represent an early stage of endoderm development. Later in development, the pancreas is formed as an endoderm-derived organ that contains insulin-secreting beta cells. We analysed whether Foxa2 binding sites in transient/stable/late or pancreatic beta cells show a different chromatin profile already before Foxa2 expression, in ESCs. As controls for active and repressed region-associated factors, we analysed binding sites of Nanog (active TF) and Trim28 (repressor). B) Left panel: Density plots showing average levels of active (H3K4me1, H3K4me3, H3K27ac, 5hmC) and repressive (H3K27me3, H3K9me3, H4K20me3, 5mC) chromatin modifications in pluripotent ES cells at specific peak sets: Nanog binding sites in ES cells, Foxa2 transient, stable, late binding sites, Foxa2 binding sites in pancreas (beta cells), Trim28 binding sites in ES cells and random genomic regions. Right panel: density plots for active and repressive chromatin modifications in d0 (black), d3F (green) and d5FS (red) cells at Nanog and Foxa2 binding sites.

We investigated by ChIP-seq active (H3K4me1, H3K4me3, H3K27ac) and repressive (H3K9me3, H3K27me3, H4K20me3) histone marks. Further, we analyzed DNA methylation (5mC) and hydroxymethylation (5hmC) by meDIP-seq. Remarkably, endoderm and beta cell-specific binding sites show a distinct chromatin signature in ESCs (Fig 6B, Fig S6A-H). Active modifications are selectively present on endoderm-specific binding sites, although at much lower levels as compared to Nanog binding sites. Beta cell-specific binding sites lack these active modifications, but also do not show prominent enrichment of repressive marks (Fig S6H, Table S7). Heatmap representations of our data demonstrate that active chromatin modifications are detectable on the majority of transient and stable Foxa2 binding sites and at a somewhat lower level on late binding sites (Fig S6D-G). During endoderm differentiation, active histone modifications on endodermal Foxa2 binding sites were further elevated, whereas no change was observed on beta cell binding sites (Fig 6B).

In summary, these data suggest that Foxa2 preferentially binds to regions of slightly active chromatin. Repressive modifications, in contrast, were largely absent in all Foxa2 binding sites (Fig 6B, Fig S6H). Notably, while re-analysing published datasets on endoderm differentiation of human ES cells (Donaghey et al., 2018; Gifford et al., 2013; Loh et al., 2014), we also observed higher levels of active chromatin marks on endoderm-specific vs. liver-specific FOXA2 binding sites (Fig S6I, Table S7), suggesting that binding preferences are evolutionarily conserved.

### Transcriptional and epigenetic effect of Foxa2 and Gata4 binding in ESCs

Our data suggest that chromatin in ESCs is prepared to favour Foxa2 binding to endoderm-specific binding sites. However, Foxa2 recruitment could be modulated by collaborating endoderm transcription factors (Donaghey et al., 2018). Thus, we wondered if Foxa2 would prefer endoderm-specific binding sites in ESCs, where endoderm TFs are not yet expressed.

We engineered doxycycline (Dox) inducible Foxa2-Venus ESCs (ESC^iFVF^), which allowed FACS isolation of Foxa2 expressing ESCs after 1, 2 and 4 days of Dox induction (Fig 7A, S7A). To test if Foxa2 expression in ESCs would already be sufficient to activate the endoderm network we performed RNA-seq of Foxa2-expressing (d2-FVFp) vs. non-expressing (d2-FVFn) ESC^iFVF^ cells 2 days after Dox induction. We found 229 differentially expressed genes (fold change > 2, padj < 0.05; Fig 7B, Table S8). Most genes were upregulated (221 up-regulated, 8 down-regulated), suggesting an activating role of Foxa2. Remarkably, only 72 out of 588 genes which were normally induced during endoderm differentiation were also upregulated in d2-FVFp cells (Table S8). Key endoderm TFs were not properly induced (Fig S7B), demonstrating that Foxa2 expression in ESCs is insufficient for endoderm differentiation.

**Figure 7.**
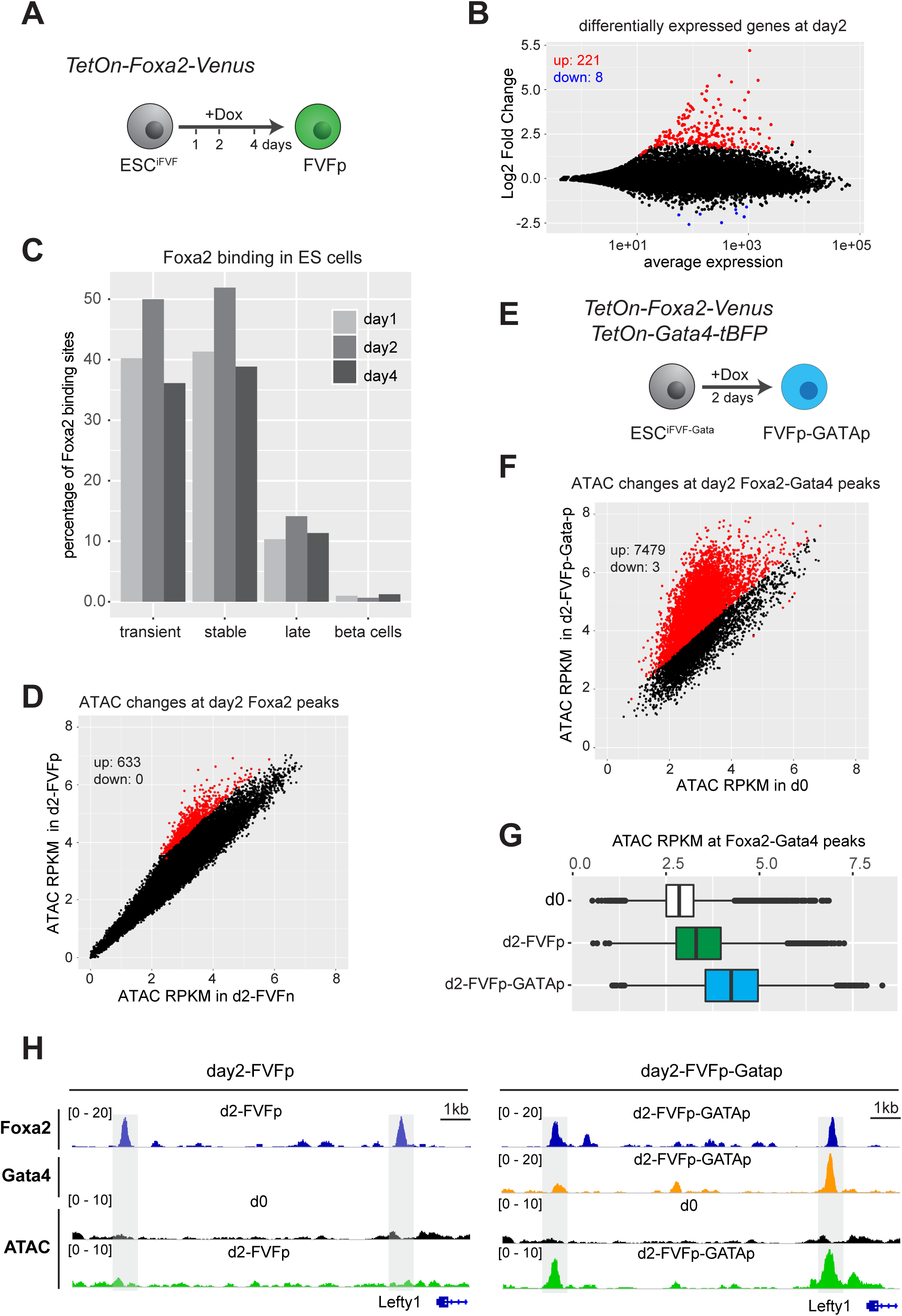
Transcriptional and epigenetic effect of Foxa2 and Gata4 binding in ES cells. A) Scheme of the experimental strategy to induce the expression of Foxa2 in ESCs. B) Dot plot showing average expression vs. log2-fold change of protein coding genes in Foxa2-expressing (d2-FVFp) vs. non-expressing (d2-FVFn) ESC^iFVF^ cells 48h after dox induction. Genes with significantly changed expression (padj < 0.05, fold change > 2; n=2 for each condition) are coloured (red = increased expression in d2-FVFp, blue = increased expression in d2-FVFn). C) Bar graph showing the percentage of beta cell- or endoderm-specific Foxa2 binding sites bound by Foxa2 in FVFp cells, one, two or four days after dox induction. D) Dot plot showing normalized ATAC-seq coverage in Foxa2-expressing (d2-FVFp) vs. non-expressing (d2-FVFn) ESC^iFVF^ cells at Foxa2 binding sites 48h after dox induction. Significant chromatin accessibility changes (padj < 0.05, fold change > 2; n=2 for each condition) are coloured in red. E) Scheme of the experimental strategy to induce the expression of Foxa2 and Gata4 in ESCs. F) Dot plot showing normalized ATAC-seq coverage in Foxa2/Gata4 co-expressing (d2-FVFp-GATAp) vs. non-expressing (d0) cells at Foxa2/Gata4 binding sites 48h after dox induction. Significant chromatin accessibility changes (padj < 0.05, fold change > 2; n=2 for each condition) are coloured in red. G) Box plots of normalized ATAC-seq coverage of Foxa2/Gata4 co-bound sites in control (d0), Foxa2-expressing (d2-FVFp) and Foxa2/Gata4 co-expressing (d2-FVFp-GATAp) cells. n=2 for each condition. Wilcoxon ranks-sum test statistics is shown in Table S7. H) Genome browser view of example Foxa2 binding sites in d2-FVFp cells (right panel, day2-FVFp) and Foxa2/Gata4 co-bound sites in d2-FVFp-GATAp cells (left panel, day2-FVFp-GATAp). The following tracks are displayed: Foxa2 Chip-seq in d2-FVFp and d2-FVFp-GATAp cells; Gata4 ChIP-seq in d2-FVFp-GATAp cells; ATACseq in d0, d2-FVFp and d2-FVFp-GATAp cells. Dashed regions indicate Foxa2 binding sites in d2-FVFp cells (left panel) which are co-bound with Gata4 in d2-FVFp-GATAp cells (right panel).

Next, we examined to which extent endodermal Foxa2 binding sites are bound by Foxa2 in ESC^iFVF^ cells 1, 2 and 4 days after Dox induction. We found that a large percentage of transient and stable peaks, but less late peaks were bound by Foxa2 in ES^FVF^ cells (Fig 7C). In contrast, only ∼1% of pancreas-specific binding sites were bound by Foxa2 in ESCs (Fig 7C). We did not observe striking changes in Foxa2 localization with longer induction times (Fig 7C). These data demonstrate that endodermal Foxa2 binding sites are primed for Foxa2 binding already in ESCs and, that Foxa2 can recognize these sites in the absence of additional endoderm-specific transcription factors. We also conclude that Foxa2 expression alone is insufficient to induce processes which would make pancreas-specific beta cell binding sites accessible.

We then performed ATAC-seq on Foxa2-expressing ESCs (d2-FVFp) to test if Foxa2 can induce chromatin accessibility on its binding sites. Remarkably, only a very small fraction (∼1%) of Foxa2 bound regions displayed significant gains in chromatin accessibility (Fig 7D), suggesting that Foxa2 binding alone is insufficient to trigger chromatin opening. We also performed ChIP-seq analyses for H3K4me1 and H3K27ac in Foxa2-expressing ESCs (d2-FVFp). We detect an increase in H3K4me1 and to lesser extent in H3K27ac (Fig S7C,D, Table S7). This behaviour mimics our findings for transient vs. stable and late binding sites during endoderm differentiation, where Foxa2 binding coincides with increased active chromatin marks but combinatorial binding with additional TFs was necessary for chromatin opening. We therefore sought for features which could distinguish binding sites showing higher chromatin accessibility vs. binding site which do not change. Firstly, we detected that ATAC-seq coverage was higher in opened Foxa2 binding sites already before Foxa2 induction (Fig S7E). Secondly, we found slightly higher enrichment for DNA binding motifs of AP1 family members in opened Foxa2 binding sites (Fig S7F). Together, our data are in line with a model in which Foxa2 preferentially binds to primed binding sites without the need for collaborating TFs. Increased chromatin accessibility, however, requires the binding of collaborating TFs in the vicinity.

To examine if collaboration between Foxa2 and additional transcription factors favours chromatin accessibility in the ES cell system, we tested whether co-expression of Foxa2 and Gata4 would result in enhanced chromatin accessibility on co-bound sites. For this experiment we generated an ESC line allowing Dox-mediated induction of both Foxa2 and Gata4 (ES^FVF-Gata^, Fig 7E). We isolated Foxa2/Gata4 double-positive cells by FACS sorting (Fig S7G) and performed ChIP-seq for Foxa2 and Gata4 as well as ATAC-seq to detect changes in chromatin accessibility. Compared to Foxa2 expressing cells (d2-FVFp), which exhibit marginally increased chromatin accessibility (Fig. 7D), Foxa2/Gata4 co-expressing cells (d2-FVFp-GATAp) showed a marked increase in chromatin accessibility on Foxa2/Gata4 co-bound sites (Fig. 7F-H). These data, together with our finding that Foxa2 and Gata4 co-binding coincides with increased chromatin accessibility during endoderm differentiation provide strong support for our hypothesis that co-binding of Foxa2 with additional TFs is needed to generate increased chromatin accessibility.

## Discussion

Pioneer transcription factors have critical roles in cell fate specification and are needed for the activation of lineage programs in a cell type-specific manner. How PTFs recognize cell type-specific target sites, and how they initiate remodelling of the surrounding chromatin remains poorly understood. In the present work we addressed these questions by studying the paradigm PTF Foxa2 in the physiological context of endoderm differentiation. Our data support a model by which Foxa2 binding sites are defined by low levels of active chromatin modifications and where local chromatin opening requires co-binding of additional transcription factors in close vicinity (Fig 8). This model is based on the following observations:

**Figure 8.**
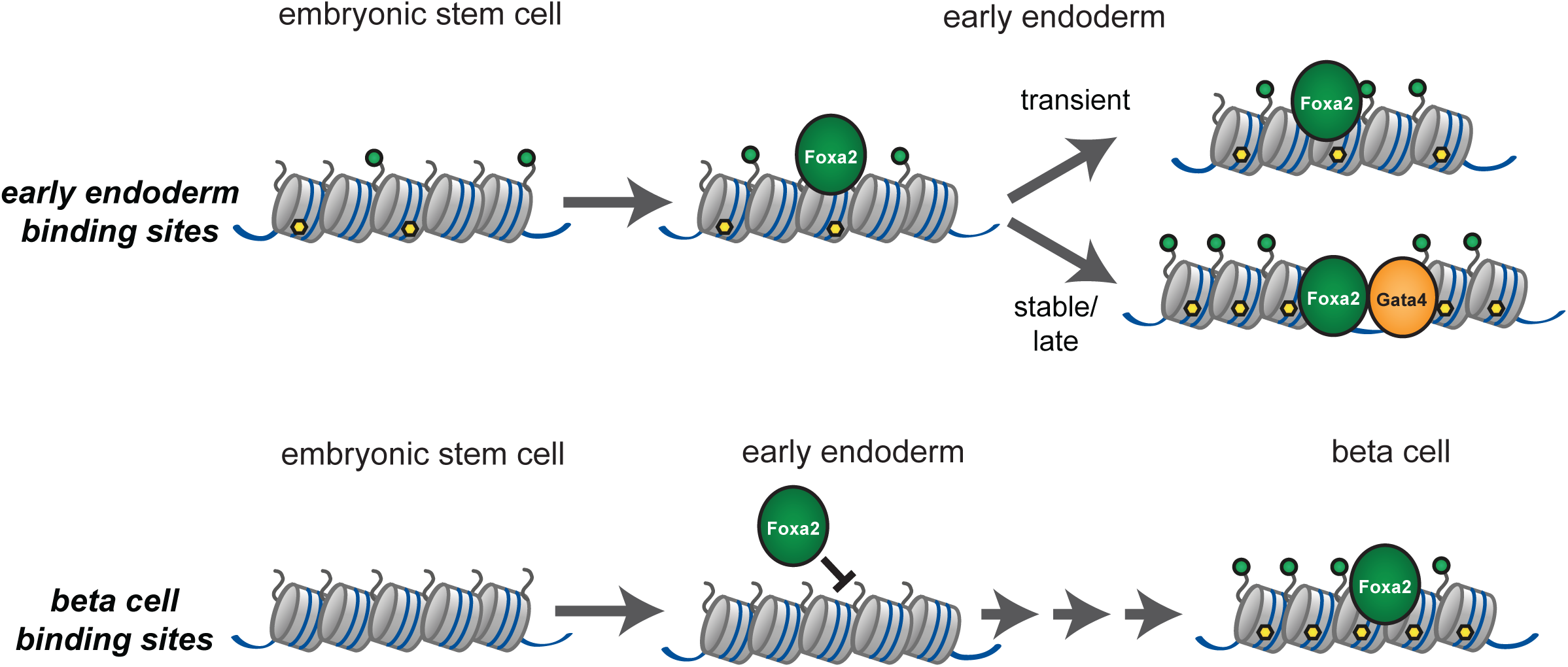
Model. Model for binding site selection and chromatin opening by Foxa2. During the transition from ESC to endoderm, Foxa2 preferentially binds endoderm binding sites featured by low levels of active chromatin modifications. Non-bound lineage-inappropriate binding sites (e.g. beta cell-specific sites) are not featured by active chromatin marks during early endoderm differentiation. Increase in chromatin accessibility occurs on binding sites where Foxa2 co-binds with additional transcription factors (stable and late sites), whereas isolated Foxa2 binding sites do not show increase in chromatin accessibility upon Foxa2 binding (transient sites).

I. In mouse ESCs, endodermal but not pancreatic Foxa2 binding sites are pre-marked by low levels of active chromatin modifications. This feature is also conserved in human ESCs.
II. Foxa2 preferentially binds to endoderm-specific, but not pancreas-specific binding sites when expressed in ESCs.
III. During endoderm differentiation, increased chromatin accessibility is observed on binding sites where Foxa2 binds together with other transcription factors, i.e. Gata4.
IV. In the absence of other endoderm transcription factors, in ESCs, Foxa2 has a very limited activity to induce chromatin opening. In the small subset of binding sites where chromatin accessibility is enhanced, Foxa2 may bind together with additional TFs, e.g. AP1 proteins.
V. Co-expression of Foxa2 and Gata4 in ESCs results in enhanced chromatin accessibility at Foxa2/Gata4 co-bound sites.

Our data are in line with recent models in which epigenetic priming determines cellular competence (Wang et al., 2015). In this way the epigenetic landscape of a cell directs transcription factor binding to lineage-appropriate sites. These regions of low level active chromatin modifications are likely to be established by the transcription factor network which is active before lineage decisions are made. In ESCs, pluripotency-associated TFs are likely to be responsible for this epigenetic priming (Kim et al., 2018).

How epigenetic priming may direct transcription factors binding is still unclear. Pioneer transcription factors, such as Foxa2, can bind specific DNA sequences in the context of nucleosomes. However, these binding sites occur very frequently in the genome. Thus, a limited number of Foxa2 molecules per cell will preferentially enrich on a set of binding sites to which Foxa2 has highest affinity. In the cellular context, affinity is not a function of DNA sequence binding alone, but rather represents a combination of different features including DNA shape, DNA methylation, chromatin organization and protein interactions in the vicinity of the binding site (Swift and Coruzzi, 2017). In this context, regions of active chromatin modifications are characterized by higher chromatin dynamics and generally enhanced accessibility, which may favor Foxa2 binding. Chromatin-modifying enzymes that reside in regions of active chromatin may also target Foxa2, thereby affecting binding affinity. In line with this hypothesis are findings that Foxa2 activity is augmented by p300-mediated acetylation on Lys259 (von Meyenn et al., 2013), whereas SIRT1-mediated deacetylation leads to reduced Foxa2 stability (van Gent et al., 2014).

From our data it is currently not possible to establish a causative link between a pre-existing chromatin state and Foxa2 recruitment. We attempted to answer this question by targeting H3K27ac to pancreas-specific Foxa2 binding sites using a Cas9-p300 fusion protein (Klann et al., 2017; Pulecio et al., 2017). Unfortunately, we were unable to detect significant levels of H3K27ac (data not shown), suggesting that establishment of a low-level active chromatin state requires more than recruitment of a single chromatin modifying factor.

Another determinant of transcription factor binding site selection could be co-binding with additional transcription factors. For example, sexual dimorphisms in liver cancer is determined by differential target activation depending on Foxa1/2 and AR or ERa interactions (Li et al., 2012b). Similarly, Oct4 occupies different genomic regions when expressed alone or in combination with other reprogramming factors (Chronis et al., 2017). Our data suggest that many Foxa2 binding sites during endoderm differentiation do not require co-binding with additional transcription factors, e.g. transient binding sites. We rather find that co-binding leads to changes in chromatin accessibility, which is likely to be a prerequisite for enhancer activation. We specifically investigated the co-binding of Foxa2 with Gata4 which occurs on stable and late binding sites. However, other endoderm-related transcription factors are likely to act in addition to Gata4 to promote chromatin opening. Interestingly, transient binding sites largely lack binding motifs except for the Foxa2 motif. The function of these binding sites is therefore rather unclear. It is possible that Foxa2 binding on transient sites is important for epigenetic priming of alternative lineages, e.g. distinct cardiac progenitors which derive from the Foxa2^+^ mesendoderm lineage (Bardot et al., 2017). However, we cannot exclude the possibility that many binding events are neutral and do not have consequences for transcriptional regulation.

Why co-binding of TFs appears necessary for inducing chromatin accessibility is still unclear. As chromatin opening is facilitated by TF-recruited chromatin remodelling activities (Swinstead et al., 2016b), co-binding of TFs could synergize in recruiting multiple chromatin remodelling machineries. Interestingly, co-binding of transcription factors in close vicinity does not immediately result in enhanced chromatin accessibility. For example, we observed on stable Foxa2 binding sites that chromatin opening is delayed. Although Foxa2 and Gata4 bind already in d3F cells, opening on these binding sites was mainly observed in d5FS cells. Delayed chromatin opening upon PTF binding was also observed recently for the PTF Pax7 in pituitary glands (Mayran et al., 2018). These findings suggest that cell cycle, replication or additional co-factors may be required for inducing higher chromatin accessibility. Experiments that specifically address the combinatorial logic of transcription factor binding and recruitment of chromatin-modifying activities are needed to better understand the requirements for enhancer activation.

## Methods

### Endoderm differentiation of DKI mESCs

DKI ESCs (Foxa2-Venus heterozygous; Sox17-Cherry homozygous) (Burtscher et al., 2013; Burtscher et al., 2012) were thawed on gamma-irradiated feeders and maintained undifferentiated in ESC medium based on DMEM (12634028, Gibco) containing 15% FCS, mLIF (self-made), 12 ml HEPES 1M (2503024, Gibco), 5 ml Penicillin/Streptomycin (15140122; Gibco), and 1 ml 2-mercaptoethanol (Gibco, 31350-010). In vitro differentiation of the ESCs towards endoderm was carried out in monolayer on 0.1% gelatine coated dishes. The cells were mouse embryo fibroblast feeder cells (MEF) depleted and cultured for few consecutive passages on gelatine and ESC medium. On the day of differentiation, ESCs were seeded (2.8 million cells for 3 days differentiation and 2.1 million cells for 5 days differentiation) on 10 cm gelatine coated dishes directly in endoderm differentiation medium (EDM) consisting of 500 ml Advanced DMEM / F-12 (1x) (Gibco/LifeTechnologies; 12634-10-500 ml), 500 ml Advanced RPMI 1640 (1x) (Gibco/LifeTechnologies; 12633-012-500 ml), 22 ml GlutaMAXTM – I CTSTM (Gibco/LifeTechnologies; 12860-01-100 ml), 200 µl AlbuMAX 100mg/ml (Gibco/LifeTechnologies; 11021-029 100g, 22 ml HEPES 1M (Gibco/LifeTechnologies; 15630-056-100 ml), 70 µl Cytidine 150 mg/ml (SIGMA; C4654-5G), 0,9 ml ß-Mercaptoethanol 50mM (Gibco/LifeTechnologies; 31350-10-20 ml), 12 ml Pen/Strep (10000U/ml) (Gibco/LifeTechnologies; 10378016 – 100 ml), 1 ml Insulin-Transferin-Selenium Ethanolamine (Gibco/LifeTechnologies; 51500-056-10 ml), supplemented with 1 ng/ml of murine Wnt3a (1324 WN-CF, R and D systems) and 10 ng/ml of Activin A (338-AC, R and D systems). Freshly prepared EDM supplemented with Wnt3a and Activin A was added every day. Cells were collected on day3 and day5 for FACS isolation and routinely tested for mycoplasma contamination.

### Endoderm differentiation of Foxa2^Venus^ ESCs

Prior to endoderm differentiation Wnt3a feeder cells (0,15×10^6/ well) (Kispert et al., 1998) were seeded on 0.1% gelatin coated 6 well plates in endoderm differentiation medium (EDM), consisting of 500 ml advanced DMEM/F-12, 500 ml advanced RPMI, 2.2x GlutaMAX, 20 mg/l Albumax, 22mM HEPES, 10 µg/ml Cytidine, 0.045mM ß-mercaptoethanol. In parallel, the Foxa2-Venus KO ESCs were split on mitomycin-treated MEFs in ESC medium without LIF. The following day, the ESCs (C59 het, C63 het, C9 homo and C17 homo) were pre-plated twice to isolate the ESCs from the mitomycin-treated feeder cells and subsequently 0,6×10^6 ESCs were seeded on the Wnt3a feeders cells in 1:1 Wnt3a conditioned medium and EDM containing Activin A (24ng/ml, final conc: 12ng/ml). 24 hours later the medium was replaced with EDM containing Activin A (12ng/ml) and refreshed every day. Cells were collected on day3 for FACS isolation and routinely tested for mycoplasma contamination.

### RNAseq of DKI cells

Total RNA from two independent biological replicates of day0, day3F+ and day5FS+ was isolated employing the RNA Clean & Concentrator kit (Zymo Research) including digestion of remaining genomic DNA according to producer’s guidelines. The Agilent 2100 Bioanalyzer was used to assess RNA quality and only high quality RNA (RIN > 8) was further processed for removal of ribosomal RNA with the Ribo-Zero Magnetic Gold Kit (Human/Mouse/Rat; Illumina). Ribosomal-depleted RNA was used as input for library preparation with Illumina TruSeq V2 RNA prep kit and processed according to the manufacturer’s instruction. Libraries were quality controlled by Qubit and Agilent DNA Bioanalyzer analysis. Deep sequencing was performed on a HiSeq 2500 system according to the standard Illumina protocol for 100bp paired end reads with v3 sequencing reagents.

### RNAseq of Foxa2^Venus^ and Doxy-Foxa2 cells

Total RNA from FACS-sorted cells was isolated employing RNA Clean & Concentrator kit (Zymo Research) including digestion of remaining genomic DNA according to producer’s guidelines. The Agilent 2100 Bioanalyzer was used to assess RNA quality and only high-quality RNA (RIN > 8) was further processed for cDNA synthesis with SMART-Seq v4 Ultra Low Input RNA Kit (Clontech cat. 634888) according to the manufacturer’s instruction. cDNA was fragmented to an average size of 200-500 bp in a Covaris S220 device (5 min; 4°C; PP 175; DF 10; CB 200). Fragmented cDNA was used as input for library preparation with MicroPlex Library Preparation Kit v2 (Diagenode, cat. C05010012) and processed according to the manufacturer’s instruction. Libraries were quality controlled by Qubit and Agilent DNA Bioanalyzer analysis. Deep sequencing was performed on a HiSeq 1500 system according to the standard Illumina protocol for 50bp single-end reads with v3 sequencing reagents.

### ChIP-seq of histone modifications

1-2 million FACS-sorted cross-linked cells (1% formaldehyde, 10min RT) were lysed in 100 ul Buffer-B (50 mM Tris-HCl, pH 8.0, 10 mM EDTA, 1%SDS, 1x protease inhibitors -Roche) and sonicated in a microtube (Covaris; 520045) using a Covaris S220 device until most of the DNA fragments were 200-500 base pairs long (settings: temperature 4°C, duty cycle 2%, peak incident power 105 Watts, cycles per burst 200). After shearing, lysates were centrifuged 10min, 4°C, 12000g and supernatant diluted with 900ul of Buffer-A (10 mM Tris-HCl, pH 7.5, 1 mM EDTA, 0.5 mM EGTA,1% Triton X-100, 0.1% SDS, 0.1% Na-deoxycholate, 140 mM NaCl, 1x protease inhibitors-Roche). 150 ul of sonicated chromatin was then incubated 4h at 4°C on a rotating wheel with 3 µg of antibody conjugated to 10 µl of Dynabeads. The antibodies used were: anti-H3K4me1 (Diagenode; Pab-037-050), H3K4me3 (Diagenode; Pab-003-050), H3K27ac (Diagenode; Pab-174-050), H3K27me3 (Diagenode; Pab-069-050), H3K9me3 (Diagenode; Pab-056-050), H4K20me3 (Diagenode; Pab-057-050). Beads were washed four times with Buffer-A (10 mM Tris-HCl, pH 7.5, 1 mM EDTA, 0.5 mM EGTA,1% Triton X-100, 0.1% SDS, 0.1% Na-deoxycholate, 140 mM NaCl, 1x protease inhibitors-Roche) and once with Buffer-C (10 mM Tris-HCl, pH 8.0, 10 mM EDTA). Beads were re-suspended in 100µl elution buffer (50 mM Tris-HCl, pH 8.0, 10 mM EDTA, 1% SDS) and incubated 20min at 65 °C. Supernatant was transferred to a new tube. Crosslink reversal of immunoprecipitated DNA was carried out overnight at 65°C. Then 100µl TE (10 mM Tris-HCl, pH 8.0, 1 mM EDTA) was added, RNA was degraded by 4μl RNase A (10mg/ml) for 1 hour at 37°C and proteins were digested with 4μl Proteinase K (10 mg/ml) at 55°C for 2 hours. Finally, DNA was isolated by phenol:chloroform:Isoamyl alcohol purification followed by ethanol precipitation. Purified DNA was used as input for library preparation with MicroPlex Library Preparation Kit v2 (Diagenode, cat. C05010012) and processed according to the manufacturer’s instruction. Libraries were quality controlled by Qubit and Agilent DNA Bioanalyzer analysis. Deep sequencing was performed on HiSeq 1500/2500 systems according to the standard Illumina protocol for 50bp single-end reads using v3 reagents.

### ChIP-seq of transcription factors

1-2 million FACS-sorted cross-linked cells (1% formaldehyde, 10min RT) were lysed in 100 ul Buffer-B-0.3 (50 mM Tris-HCl, pH 8.0, 10 mM EDTA, 0,3%SDS, 1x protease inhibitors -Roche) and sonicated in a microtube (Covaris; 520045) using a Covaris S220 device until most of the DNA fragments were 200-500 base pairs long (settings: temperature 4°C, duty cycle 2%, peak incident power 105 Watts, cycles per burst 200). After shearing, lysates were centrifuged 10min, 4°C, 12000g and supernatant diluted with 1 volume of Dilution Buffer (1mM EGTA 300 mM NaCl, 2% Triton x-100, 0.2% sodium deoxycholate, 1x protease inhibitors-Roche). Sonicated chromatin was then incubated 4h at 4°C on a rotating wheel with 6 ug of antibody conjugated to 20 µl of Dynabeads. The antibodies used were: anti-Foxa2 (SantaCruz; sc6554x), anti-Gata4 (R&D Systems; AF2606), anti-Nanog (Bethyl lab; A300-397-A). Beads were washed four times with Buffer-A (10 mM Tris-HCl, pH 7.5, 1 mM EDTA, 0.5 mM EGTA,1% Triton X-100, 0.1% SDS, 0.1% Na-deoxycholate, 140 mM NaCl, 1x protease inhibitors) and once with Buffer-C (10 mM Tris-HCl, pH 8.0, 10 mM EDTA). Beads were then incubated with 70 μl elution buffer (0.5% SDS, 300 mM NaCl, 5 mM EDTA, 10 mM Tris HCl pH 8.0) containing 2 μl of Proteinase K (20mg/ml) for 1 hour at 55°C and 8 hours at 65°C to revert formaldehyde crosslinking, and supernatant was transferred to a new tube. Another 30 μl of elution buffer was added to the beads for 1 minute and eluates were combined and incubated with another 1 μl of Proteinase K for 1 hour at 55°C. Finally, DNA was purified with SPRI AMPure XP beads (Beckman Coulter) (sample-to-beads ratio 1:2). Purified DNA was used as input for library preparation with MicroPlex Library Preparation Kit v2 (Diagenode, cat. C05010012) and processed according to the manufacturer’s instruction. Libraries were quality controlled by Qubit and Agilent DNA Bioanalyzer analysis. Deep sequencing was performed on HiSeq 1500/2500 systems according to the standard Illumina protocol for 50bp single-end reads using v3 reagents.

### meDIP-seq and Hydroxy-meDIP-seq

The procedure was adapted from (Maunakea et al., 2010; Mohn et al., 2009). Genomic DNA from FACS-sorted cells was randomly sheared to 100-500 bp in a microtube (Covaris; 520045) using a Covaris S220 device (400 sec; 4°C; PP 140; DF 10; CB 200). Sonicated DNA was end-repaired, A-tailed, and ligated to Illumina multiplex adaptors according to NEBNext DNA library prep kit (NEB E6040S). Ligated DNA was purified using Agencourt AMPure XP beads (Beckman Coulter). 1 µg of adaptor-ligated DNA was used for each immunoprecipitation and heat-denatured at 95°C for 10min, rapidly cooled on ice and immunoprecipitated overnight at 4 °C with rocking agitation in 500 ml immunoprecipitation buffer (10mM sodium phosphate buffer, pH 7.0, 140mM NaCl, 0.05% Triton X-100) using 1 µl of mouse monoclonal anti-5-methylcytosine antibody (Eurogentec BI-MECY-0100) or 0,5 µl of rabbit 5-Hydroxymethylcytosine antibody (Active motif 39769). To recover the antibody-bound DNA fragments, 50µl Protein G Dynabeads (ThermoFisher) and, only to anti-5-methylcytosine IPs, 5µl of rabbit anti-mouse IgG secondary antibody (Active Motif cat.53017) were added and incubated for an additional 2 h at 4 °C with agitation. After immunoprecipitation a total of seven to ten immunoprecipitation washes were performed with ice-cold immunoprecipitation buffer. Washed beads were resuspended in TE buffer with 0.25% SDS and 0.25 mg/ml proteinase K for 2 h at 55 °C with vigorous shaking (900 rpm). DNA was purified with the Qiagen PCR clean-up MinElute kit (Qiagen) and eluted in 30 ul. Samples were then amplified by PCR with Illumina primers (NEBNext Multiplex Oligos for Illumina cat. E7335) in a 50⍰ μl reaction with 2× PCR master mix (NEB cat. M0541). PCR cycled as: (1) 98⍰ °C, 30s; (2) 98⍰ °C, 10⍰ s; (3) 60⍰ °C, 30⍰ s; (4) 72⍰ °C, 30s; (5) repeat steps (2)–(4) for 4-10 cycles; (6) 72°C, 5min. Amplified libraries were purified using Agencourt AMPure XP beads (Beckman Coulter). Quality control was carried out with a Qubit fluorometer and a Bioanalyzer (Agilent). 50 bp single-end sequencing was performed with a a HiSeq 1500 sequencer with v3 reagents (Illumina).

### ATAC-seq

ATAC-seq was done as previously described (Buenrostro et al., 2013). Briefly, 50000 FACS sorted cells were washed in 1xPBS, re-suspended in 50 ul of lysis buffer (10 mM Tris pH 7.4, 10 mM NaCl, 3 mM MgCl2, 0.1% NP40,) and spun at 500g for 10 min at 4 °C to collect nuclei. Nuclei were subsequently re-suspended in 50 µl Transposase reaction containing 25µL 2x tagmentation buffer, 22.5 µl water, 2.5 µl Tn5 Transposase (Illumina Nextera DNA Library Preparation Kit, cat. FC-121-1030). Reactions were incubated for 30 min at 37° C in a thermomixer shaking at 300 rpm and DNA purified using Qiagen PCR clean-up MinElute kit (Qiagen). The transposed DNA was subsequently amplified in 50µl reactions with custom primers as described (Buenrostro et al., 2013). After 4 cycles libraries were then monitored with qPCR: 5 µl PCR sample in a 15⍰µl reaction with the same primers. qPCR output was monitored for the ΔRN; 0.25 ΔRN cycle number was used to estimate the number of additional cycles of the PCR reaction needed for the remaining PCR samples. Amplified libraries were purified with the Qiagen PCR clean-up MinElute kit (Qiagen) and size selected for fragments less than 600 bp using the Agencourt AMPure XP beads (Beckman Coulter). Libraries were quality controlled by Qubit and Agilent DNA Bioanalyzer analysis. Deep sequencing was performed on a HiSeq 1500 system according to the standard Illumina protocol for 50bp single-end reads or paired-end reads.

### Fluorescence Activated Cell Sorting

For RNAseq, ATACseq and meDIP-seq, following trypsin treatment, cells were resuspended in PBS with 10% FCS before FACS collection. For ChIP-seq cells, cells were fixed for 10min with 1% Formaldehyde and quenched with 0.125 M final concentration glycine before FACS collection. FACS was performed with a FACSAria instrument (BD Biosciences). Data were analyzed with FlowJo software.

### Generation of Foxa2^Venus^ Knockout cells

#### Generation of Foxa2^Venus^ knockout targeting vector

The knockout construct was designed as shown in Fig.S3. The pBKS-Venus vector carrying the H2B-Venus sequence (Nagai et al., 2002) was linearized with SalI. To induce a blunt end and a NotI restriction site the 5’ overhangs of the SalI linearized vector were filled up using Klenow enzyme and subsequently cut with NotI. The 3’ homology arm for the Foxa2 gene was cloned from the pBKS-3’HA vector into the linearized pBKS-H2B-Venus vector using the restriction enzymes EcoRV and KpnI, resulting in pBKS-Venus-3’HA. The 5’ homology arm for the Foxa2 gene was amplified from the pL254-Foxa2 vector (Burtscher et al., 2013) as a template using primers carrying a NotI site (EP 1447/EP 1448). The amplified 5’ homology arm was cloned via NotI into the pBKS-Venus-3’HA vector, resulting in p-BKS-Venus-3’HA-5’HA vector, containing the H2B-Venus sequence flanked by the 5’ and 3’ homology arms of the Foxa2 gene. In between the H2B-Venus and 3’ homology arm the SV40 polyadenylation signal sequence followed by the loxP-flanked murine PGK and the bacterial EM7 promoter-driven neomycin resistance gene. To ensure the correct insertion of the inserts, the vectors were digested with HindIII/XhoI: pBKS-Venus: 4574bp, 1910bp; pBKS-Venus-3’HA: 4574bp, 1912bp, 484bp; and with ApaLI/XhoI: p-BKS-Venus-3’HA-5’HA: 1246bp, 928bp, 727bp, 602bp wrong orientation: 1246bp, 1206bp, 727bp, 602bp and pBKS-Venus-3’HA: 2601bp, 2396bp, 1246bp, 727bp. Subsequently, the Foxa2-Venus targeting vector was sequenced and the vector with the correct sequence was used for transfection of the mESCs. Prior to transfection of mESCs the targeting vector was linearized using ApaI.

#### Oligonucleotides used for genotyping and sequencing

EP 1447 NNNGCGGCCGCGGGAATGTGCACTAAAAGGGAGGAAACC

EP 1448 NNNGCGGCCGCGCATACTGGAAGCCGAGTGCATGG

EP 397 CTACTACCAAGGAGTGTACTCC

EP 1520 ACCATTACGCCTTCAACCAC

EP 1513 GGAATTCTGGCCATTCTAGC

EP 1499 GGCTGGACGTAAACTCCTCTTC

#### Foxa2 3’ Southern probe (732bp)

CTGGATATGCTCTAGAAAGGCAGAAGTTTACAGTTTTTTTAATATCAGGCCTCCTTTCTAGTCAGTGAACTTAGACTGGGTTTACCAATTTTGGTGCATGGCTCTTCCAGCTACTTGAAGCATTGCCCCCCCTAGACCTTCCTGTGCCATTGAGACTACCTGGCTCTAGGTTGTGCCGGGAGGGCAGCCTGTCTCAGTCTCACAGGTGTTATCCAGGTATTGGGAAACCTTGCTAGGCTAGGAACGATGAGCCACCTAATCTGGGGAAACATTTTAACATTGGGAATTGGGTATAATTGCATAGTTAAGGGTAACCCCCAAATCTTTTATTAAGAAGTTATTCTGTGGGTGGGGAGATAGGGAGGGATGGAAGGGTGCCCTGAGCAGCTTAGCAAATGACTCCCAAAGTAGTGAAATCCCAGTGTCTCAGGAATGGTGTCTCCCTTCTACCAGCCAGGGCAAAGCTGTTTGTTAGCTTAGGAAGCTCCTATAGGCAAACCACACTTGAGGCCCAGGGACTGAATGGGTATTTTGTGAGCCTCCAGGAAAATACAAAGACCCCAAATAAAACCTCACCAATCATTTCCACCACTCTGCAGATTTTCCAAATTGACGGGTAACTGTAGAGGAGGTCGTGTTTTGCAAAAGGAGCCTCCTCACGCTGACCTGCATCTCCTGCCCTTGAAGCTGTCCCTCCCGCCCGCCCCCAGTCTGACTTTCCATAGGCCATTC

#### Design of guide RNAs (sgRNAs)

To design the gRNAs the online tool Optimized CRISPR Design (crispr.mit.edu/) was used. 250 nucleotides of the coding region in exon 3 were submitted as template for the gRNA design. The selected gRNA pair had a 52 bp 5’ overhang and 17 bp gRNA offset.

#### The submitted sequence to Optimized CRISPR Design

CTCTATCAACAACCTCATGTCGTCCGAGCAGCAACATCACCACAGCCACCACCACCATCAGCCCCACAAAATGGACCTCAAGGCCTACGAACAGGTCATGCACTACCCAGGGGGCTATGGTTCCCCCATGCCAGGCAGCTTGGCCATGGGCCCAGTCACGAACAAAGCGGGCCTGGATGCCTCGCCCCTGGCTGCAGACACTTCCTACTACCAAGGAGTGTACTCCAGGCCTATTATGAACTCATCCTAA

#### The selected gRNAs

gRNA #4 forward 5’-3’: CACCGGGGGATGAGTTCATAATAGGCC

gRNA#4 reverse 3’-5’: CCCCCTACTCAAGTATTATCCGGCAAA

gRNA#12 forward 5’-3’: CACCGGGGCCCTGCTAGCTCTGGTCAC

gRNA#12 reverse 3’-5’: CCCCGGGACGATCGAGACCAGTGCAAA

#### Cell Culture and genome targeting of mESCs via CRISPR-Cas9D10a

Mouse embryonic stem cells (mESCs) were cultured on MMC-treated murine embryonic feeders (MEFs) in Dulbecco’s Modified Eagle Medium (DMEM, Invitrogen) containing 15% fetal calf serum (FCS, PAN), 2 mM L-glutamine (Invitro-gen), 100 lM b-mercaptoethanol (Invitrogen) and 1,500U/ml LIF (self-made). Prior to electroporation, the Foxa2-Venus targeting vector was linearized via ApaI and subsequently together with the gRNAs and Cas9D10A cleaned up by phenol-chloroform extraction followed by EtOH precipitation. For each electroporation half a 10 cm dish of confluent IDG3.2 mESCs (Hitz et al., 2007) was used. The medium was refreshed the next day. After 24 hours the selection of the cells was initiated with 300 µg/ml geneticin (G418).

### Generation of Doxycycline-inducible Foxa2-Venus ES cells

We made use of a lentiviral Tet-On-3G system that consists of two lentiviral vectors: a regulator vector that stably expresses the Tet-On 3G transactivator protein (1702_pLenti6-EF1a-Tet3G-IRES-Neo), and a response vector that contains the TRE3G promoter that controls the expression of Foxa2-Venus (1701_pLenti6-TRE3G-Foxa2-Venus-PGK-PuroR).

For the 1702_pLenti6-EF1a-Tet3G-IRES-Neo, a Human EF1a Promoter, a Tet-On 3G element and a Neo resistance gene were cloned into pLenti6-puro backbone (Sadic et al., 2015) using Gibson Assembly as described (Gibson et al., 2009). The Human EF1a Promoter/Tet-On 3G fragment as well as the Neo(R) cDNA were PCR amplified from the pEF1α-Tet3G (Cat. # 631336, Clontech) using the following primer pairs

F: gggacagcagagatccactttggccgcggctcgaggagtaattcatacaaaaggactcg,

R: ccgatcgatagatcttcatgtctggatccttacttagttacc

and

F: tgccttgtaagtcattggtcttaaaggtaccctcagaagaactcgtcaagaag,

R: accgggccggatatcatgattgaacaagatggattgc, respectively. The IRES2 fragment was PCR amplified from the pTRE3G-IRES (Cat. # 631174, Clontech) using the primer pairs

F: aggatccagacatgaagatctatcgatcggccg,

R: atcttgttcaatcatgatatccggcccgg. T

he pLenti6 backbone was cut with XhoI and KpnI.

The 1701_pLenti6-TRE3G-Foxa2-Venus-PGK-PuroR was cloned in two steps using Gibson Assembly. First, the Foxa2 cDNA was amplified from the pHD-Foxa2 vector (Kindly provided by Klaus Kaestner) using the following primer pairs

F:tttccgtaccacttcctaccctcgtaaagtcgacaccggggcccagatctATGCTGGGAGCCGTGAAGATGGAAGGG

R: GGTGAACAGCTCCTCGCCCTTGCTCACCATTCTAGAGGATGAGTTCATAATAGGCCTGGAGT

and the Venus cDNA was amplified using the following primer pairs

F: GAGTGTACTCCAGGCCTATTATGAACTCATCCTCTAGAATGGTGAGCAAGGGCGAGGAGCTG,

R: agaatttcgtcatcgctgaatacagttacattggatccctgcaggctagcTTACTTGTACAGCTCGTCCATGCC.

Both fragments were then assembled together with a pTRE3G-IRES (Cat. # 631174, Clontech) backbone, cut with BglII and NheI. From the resulting vector, the TRE3G-Foxa2-Venus cassette was amplified using the following primer pair

F: tttattacagggacagcagagatccactttggccgcggctaggcgtatcacgaggccctt,

R: ctgccttggaaaaggcgcaaccccaacccccggatccctgcaggctagc and assembled together with a pLenti6-puro backbone, cut with XhoI.

Lentiviral particles were generated using standard protocols. After transduction SCF-ES cells (Sox17-Cherry homozygous, derived from the same batch of blastocysts as DKI cells) were selected with Puromycin (1ug/ml) and Neomycin (250ug/ul). Foxa2 expression was induced in cell lines T119 and T128 with 40ng/ul doxycycline for 1 day, 2 days or 4 days followed by FACS isolation. Cells were routinely tested for mycoplasma contamination.

### Generation of Doxycycline-inducible Foxa2-Venus, Gata4-tagBFP ES cells

We made use of the Tet-On-3G system already present in the Doxycycline-inducible Foxa2-Venus ES cells (cell line T128).

A lentiviral vector containing the TRE3G promoter controlling the expression of Gata4 (1722_pLV[Tet]-Bsd-TRE3G-mGata4-T2A-TagBFP) was generated by VectorBuilder (Vector ID is VB180822-1113dwn which can be used to retrieve detailed information about the vector on vectorbuilder.com).

Lentiviral particles were generated using standard protocols. After transduction the T128 cells were selected with Blasticidin (5ug/ml). Foxa2 and Gata4 expression were induced in cell line T134 with 40ng/ul doxycycline for 2 days followed by FACS isolation. Cells were routinely tested for mycoplasma contamination.

### Immunohistochemistry

The Foxa2^Venus^ mESCs were differentiated under endoderm conditions on IbiTreat µ-Slide 8 well chambers for 3 days. The mESCs were fixed in 4% PFA in DPBS for 5 min at RT, washed once and permeabilized for 10 min at RT using 0.1 M glycine and 0.1 % triton X - 100 in MilliQ water. Followed permeabilization, the cells were rinsed 2 x with DPBST and blocked in DPBST containing 0.1 % Tween - 20, 10 % heat inactivated fetal calf serum (FCS), 0.1 % BSA and 3 % donkey serum for 30 minutes at 37°C. Subsequently, the blocking solution was replaced by primary antibodies diluted in blocking solution: α-Foxa2 1:1000 (Cell signaling #8186) and α-GFP 1:1000 (Aves Labs #1020), α-Oct4 1:500 (Santa Cruz #5279), α-Sox17 1:500 (Acris/Novus #GT15094), α-T 1:300 (Santa Cruz #17743), α-Cer1 1:500 (R&D #AF1075) and incubated for 3 hours at RT while shaking. The cells were washed three times for 5 minutes each and afterwards incubated for 1 hour at RT in blocking solution containing the following secondary antibodies (1:800): α- chicken (Dianova #703-225-155) and α-rabbit (Invitrogen #A21206), α-mouse (Invitrogen #A31570), α-goat (Invitrogen #A21432). The cells were stained with DAPI 1:500 and washed three times with DPBST before taking pictures with a Leica SP5 confocal microscope.

### Quantification and statistical analysis

#### RNA-seq

Paired end or single end reads were aligned to the mouse genome version mm10 using STAR (Dobin et al., 2013) with default options “--runThreadN 32 --quantMode TranscriptomeSAM GeneCounts --outSAMtype BAM SortedByCoordinate”. Read counts for all genes were normalized using DESeq2 (Love et al., 2014). Significantly changed genes were determined through pairwise comparisons using the DESeq2 results function (log_2_ fold change threshold=1, adjusted p-value <0.05). For the endoderm differentiation time course analysis, pairwise comparisons between all states (d0, d3F, d5FS) were performed with Deseq2 to isolate a list of differentially expressed genes during endoderm differentiation. Heatmap with differentially expressed genes was plotted with pheatmap using rlog-normalized expression values. PCA analyses were done using the plotPCA function of the DESeq2 package. Bargraphs showing expression data for selected genes were plotted using ggplot2 with RSEM-normalized data (TPM = Transcript Per Million).

#### ATAC-seq

ATAC-seq reads were aligned to the mouse genome mm10 using Bowtie (Langmead, 2010) with options “-q -n 2 --best --chunkmbs 2000 -p 32 -S”. ATAC peaks over Input background were identified using Homer (Heinz et al., 2010) findPeaks.pl with option “-style factor”. Peaks from all samples were merged using mergePeaks resulting in a unified Peak set. The peak list was filtered for promoter-associate peaks (distance to TSS < 1000bp) with bedtools. Raw ATAC coverage counts were then calculated with annotatePeaks within 500bp across the peak centers. Differential ATAC peaks were determined with the DESeq2 result function and filtered for adjusted p-value < 0.05 and log2 fold change > 1. Genomic feature annotation of ATAC-seq peaks was done using annotatePeaks. Transcription factor motif prediction was done with findMotifsGenome.

ATAC coverage on Foxa2 peaks was calculated using homer annotatePeaks from replicate experiments.

ATAC-seq quality control was performed by investigating the nucleosomal pattern of the bioanalyzer profile of the ATAC-seq libraries, the number of mapped reads and the fraction of reads in peaks vs mapped reads (Supplementary Table S9).

#### ChIP-seq

ChIP-seq reads were aligned to the mouse genome mm10 using Bowtie with options “-q -n 2 --best --chunkmbs 2000 -p 32 -S”. Transcription factor peaks vs Input background were identified using findPeaks with option “-style factor”. To define specific peaks for d3F and d5FS stages, Foxa2 peaks which were shared between replicates and showed strong enrichment over input (fc>10) were identified and sorted into d3F, d3F+5FS and d5FS using the homer mergePeaks tool.

For the overlap analysis of Foxa2 binding in ES cells, endoderm and beta cells, high-confidence binding sites were defined with fold change over input > 5 and stage-specific binding sites were counter selected with fold change over input < 2.

Annotation of genomic regions was based on homer annotatePeaks. Transcription factor motif analysis was done with findMotifsGenome. GREAT analysis was done using the online resource at http://great.stanford.edu/public/html.

Coverage of histone modifications was calculated with annotatePeaks from corresponding Tag Directories. Replicate experiments were merged. Boxplots were generated with ggplot2.

Density plots for histone modifications were based on high confidence non-promoter peaks of Foxa2 in d3F and beta cells, as well as Nanog and Trim28 binding sites. Coverage density was calculated with annotatePeaks using option “-size 5000 -hist 50” from corresponding tag directories. Replicate experiments were averaged.

#### TF binding site predictions

TF binding site plots across Foxa2 binding sites were generated using annotatePeaks with option -m to detect TF motif occurrence within 5000bp of each Foxa2 binding site. Density plots were then generated using ggplot2.

#### Ranking of transcription factors

Putative lead TFs were assessed by adopting a previously published approach (Rackham et al., 2016). For each transcription factor we established a sphere of influence of up to three level depth using gene-gene relations based on the STRING database version 10.5 (Szklarczyk et al., 2017). We considered only relations with total scores >300, where less than half of the value was attributed to text mining. Using the measured difference in RNA-seq expression we calculated scores, one for the TF and one for its underling network using the following equations:

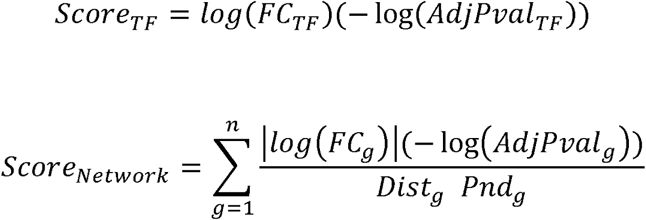

where: TF – transcription factor, FC – fold change, AdjPval – adjusted p-value (both from DSeq2), g – gene, Dist – number of steps between g and TF, Pnd – parent node degree, n - length of list of genes associated with TF.

Subsequently all factors were ranked based on combined ranking of both scores and TFs with expression values below 2 TPM at posterior stage were removed. The TFs were then plotted as network using cytoscape 3.6 (Shannon et al., 2003) with edges width correlating to the STRING interaction score. Only factors connected to other TFs were plotted and the top 5 ranked TFs were highlighted by increase node size.

### Ontology Annotation

In silico functional annotation of different groups of Foxa2 binding sites and dynamic ATAC regions were performed with the GREAT tool (McLean et al., 2010), using the default settings and the whole mouse genome as background. The terms belonging to various gene annotations (GO Biological Process, MSigDB Pathway) were considered. Differentially regulated genes defined by DESeq2 were assigned to GO biological process groups (Annotation data set: GO biological process complete release 20181115) using the PANTHER Overrepresentation Test (release 20181113) (Mi et al., 2017) with default parameters and with all mouse genes as background.

### Data and code availability

All genomic data have been deposited in the GEO database under accession number GSE116262 and will be made public upon acceptance. All the genomic data analysed in this study are listed in Table S10.

The code underlying our analysis is available upon request.

### Supplemental Data

Supplementary figures related to Figure 1, Figure 2, Figure 3, Figure 4, Figure 5, Figure 6, Figure 7.

Supplementary Tables: S1, S2, S3, S4, S5, S6, S7, S8, S9, S10.

## Acknowledgements

High throughput sequencing was performed by the Laboratory for Functional Genome Analysis (LAFUGA) of the Ludwig-Maximilian-University, Munich. Additional sequencing was done by the Norwegian Sequencing Centre (www.sequencing.uio.no), a national technology platform hosted by the University of Oslo and Oslo University Hospital, and supported by the “Functional Genomics” and “Infrastructure” programs of the Research Council of Norway and the Southeastern Regional Health Authorities.

We acknowledge the Core Facility Flow Cytometry at the Biomedical Center, Ludwig-Maxmilians-Universität München, for providing equipment, service and expertise. We also thank Maryam Kazerani for precious technical assistance in FACS sorting. We further acknowledge Christopher Breunig, Maximilian Wiesbeck and Stefan Stricker for help with dCas9-p300 mediated epigenome engineering.

For financial support we would like to thank the Helmholtz Society and German Research Foundation and German Center for Diabetes Research (DZD e.V.). Work in the G.S. lab is funded by the German Research Foundation (SPP1356; SFB1064 – A11, SFB1243 – A03, SFB1321 - P13)

## Author contributions

FMC, HL, GS conceived and designed the project; FMC performed NGS experiments; SH,ZW,KS,SG performed cell differentiation and Fluorescence Activated Cell Sorting; KS performed immunofluorescence cell staining; KS, IB generated the Foxa2^Venus^ KO ES cell line; MS generated the TenON-Foxa2-Venus lentiviral vectors; IE, GDG contributed to library preparation and NGS sequencing; PS generated the TF networks; FMC, HL, GS analysed and interpreted the results; GS performed the bioinformatic analysis; FMC and GS wrote the manuscript with the help of HL and inputs from the other authors.

## Competing interests

The authors declare that they have no competing interests.

