## Supplementary figures for "Pre-marked chromatin and transcription factor co-binding shape the pioneering activity of Foxa2"

Supplementary figures related to: Figure 1, Figure 2, Figure 3, Figure 4, Figure 5, Figure 6, Figure 7.

Supplementary tables:

Table S1_significant_genes_d0_d3F_d5FS.xlsx

Table S2_DARs_gene.association.xlsx

Table S3_motifs_ATAC-DARs.xlsx

Table S4_significant_genes_het_homo.xlsx

Table S5_Foxa2_peaks.xlsx

Table S6_motifs_Foxa2-binding-sites.xlsx

Table S7_wilcoxon.statistics.Boxplots.xlsx

Table S8_Foxa2 expression ESC - regulated_genes.xlsx

Table S9_ATACseq.metrics.xlsx

Table S10_NGS-Sample_list.xlsx

**Supplement related to Figure 1**

A) Representative FACS plots for pluripotent and differentiated ESCs at day0, day3 and day5 of *in vitro* endoderm differentiation. Differentiated cells were sorted based on expression of Foxa2 and Sox17.

B) Principal component analysis of gene expression profiles determined by RNA-seq from FACS sorted d0, d3F and d5FS cells. Dots of the same colour represent biological replicates.

C) Gene Ontology analysis showing selected biological processes enriched in genes differentially expressed in the transitions d0-d3F and d0-d5FS.

**Supplement related to Figure 2**

A) Percentages of all ATAC-seq peaks associated with different genomic features in FACS sorted d0, d3F and d5FS cells.

B) Table showing the percentages of differentially expressed genes (UP and DOWN in the indicated comparison) linked to dynamic accessible regions.

C) Principal component analysis of all ATAC-seq peaks, promoter ATAC peaks or non-promoter ATAC peaks detected in FACS sorted d0, d3F and d5FS cells. Dots of the same colour represent biological replicates.

D) Gene ontology analysis showing enriched biological processes at differentially accessible regions d0 vs. d3F.

E) Gene ontology analysis showing enriched biological processes at differentially accessible regions d0 vs. d5FS.

F) Gene ontology analysis showing enriched biological processes at differentially accessible regions d3F vs. d5FS.

**Supplement related to Figure 3**

A) Targeting strategy for the Foxa2H2B-V allele. The whole open reading frame of Foxa2 was replaced by the H2B-Venus sequence followed by the SV40 poly-A signal sequence and the loxP-flanked murine PGK promoter-driven neomycin resistance gene. The construct is flanked by the 3' and 5' homology arms for Foxa2. The 3' and 5' UTRs of Foxa2 are illustrated by black boxes, the predicted promoter regions by orange boxes and the coding region by red boxes. The Cas9D10A cleaving site is depicted as a red arrowhead. Primer binding sites are indicated by horizontal arrows and primers were used for genotyping PCR (EP 397, EP 1499, EP 1513) and indel PCR (EP 1520, EP 1513). The location of the 3' and 5' southern probes and the restriction sites are shown.

B) The mESC clones were genotyped with the primers EP 1513, EP 397 for the WT allele (649bp) and EP 1513, EP 1499 for the targeted allele (740bp). Out of 24 clones 3 clones were homozygous (12,5%), 7 clones were heterozygous (29%) and 14 clones were WT (58,5%). Indel mutations were detected by the primers EP 1513 and EP 1520 flanking the gRNA binding sites resulting in an 878bp DNA amplicon, which was sequenced. Representative sequences of the Foxa2 locus targeted by Cas9D10A are shown with gRNAs 4/12 target sites and PAMs are indicated by blue and red font color, respectively. The Cas9D10A cleaving site is illustrated by a red arrowhead. Representative indels are shown in selected sequences. Out of 7 heterozygous clones 2 clones showed the WT sequence (WT, 28,6%), whereas 4 clones had deletions (Del, 57%) and one clone had insertions (Ins, 14,4%).

C) Southern blot of mESCs digested with HindIII and hybridized with the 3' southern probe showing the 20012bp WT allele and 6800bp targeted allele. The first Foxa2 (V/V) clone shows residual WT allele from the MEFs used for mESC maintenance.

D) Confocal sections showing differentiated Foxa2 (V/V) and Foxa2 (V/+) mESCs at day 3 of endoderm differentiation stained with antibodies against GFP (green), DAPI (blue), Foxa2 (red). Scale bar: 10µm. Del: deletion, Ins: insertion, MEFs: mouse embryonic fibroblasts.

E) Representative FACS plots for Foxa2$^{Venus/+}$ undifferentiated cells (d0$^{con}$), Foxa2$^{Venus/+}$ day3 (d3$^{con}$) and Foxa2$^{Venus/Venus}$ day3 (d3$^{KO}$) endoderm differentiating cells. The gates indicate Venus positive endoderm differentiating cells.

F) Principal component analysis of gene expression profiles obtained by RNA-seq from FACS sorted pluripotent control ($d0^{con}$) and Foxa2 ko ($d0^{ko}$) cells and endoderm differentiating control ($d3^{con}$) and Foxa2 ko ($d3^{ko}$) cells. Dots of the same colour represent biological replicates.

G) Gene Ontology analysis showing the top 10 (ranked by their p-value) enriched biological processes in genes downregulated or upregulated in endoderm differentiating Foxa2 ko cells ($Foxa2^{Venus/Venus}$).

H) Average expression levels of pluripotency markers detected by RNA-seq in pluripotent control ($d0^{con}$) and endoderm differentiating control ($d3^{con}$) and Foxa2 ko ($d3^{ko}$) cells. TPM: Transcripts Per Kilobase Million. Error bars depict standard deviation (n=2 for d0, n=3 for d3).

I) Average expression levels of mesoderm markers detected by RNA-seq in pluripotent control ($d0^{con}$) and endoderm differentiating control ($d3^{con}$) and Foxa2 ko ($d3^{ko}$) cells. TPM: Transcripts Per Kilobase Million. Error bars depict standard deviation (n=2 for d0, n=3 for d3).

L) Average expression levels of endoderm markers detected by RNA-seq in day0 control ($d0^{con}$), day3 control ($d3^{con}$) and day3 ko ($d3^{ko}$) cells. TPM: Transcripts Per Kilobase Million. Error bars depict standard deviation (n=2 for d0, n=3 for d3).

**Supplement related to Figure 4**

A) Motif analysis in Foxa2 binding sites. Homer findMotifsGenome was used to identify enriched transcription factor binding motifs within 200 bp of Foxa2 peak summits. The top-scoring motif was in all cases representative of the Foxa family.

B) Gene ontology analysis for biological processes enriched at transient, stable and late Foxa2 binding sites.

C) Enrichment analysis for Molecular Signatures Database (MSigDB) pathways at transient, stable and late Foxa2 binding sites. Red and blue arrows indicate Wnt-signaling and Foxa network related pathways, respectively.

D) Bar plot showing the fraction of differentially expressed genes in d0 vs. d5FS cells bound by Foxa2.

E) Bar plot showing the fraction of differentially expressed genes in control vs. Foxa2 ko cells bound by Foxa2.

F) Scatter plots showing normalized ATAC-seq signals at Foxa2 transient binding sites. From left to right: ATAC-seq signal in d0 vs. d3F, d0 vs. d5FS, control vs. Foxa2 ko cells. Significant chromatin accessibility changes (padj < 0.05, log2 fold change > 1) are coloured in red.

G) Scatter plots showing normalized ATAC-seq signals at Foxa2 stable binding sites. From left to right: ATAC-seq signal in d0 vs. d3F, d0 vs. d5FS, control vs. Foxa2 ko cells. Significant chromatin accessibility changes (padj < 0.05, log2 fold change > 1) are coloured in red.

H) Scatter plots showing normalized ATAC-seq signals at Foxa2 late binding sites. From left to right: ATAC-seq signal in d0 vs. d3F, d0 vs. d5FS, control vs. Foxa2 ko cells. Significant chromatin accessibility changes (padj < 0.05, log2 fold change > 1) are coloured in red.

I) Genome browser view of example transient, stable and late Foxa2 binding sites. The following tracks are displayed: Foxa2 ChIP-seq in d3F and d5FS cells; ATAC-seq in d0, d3F, d5FS, $Foxa2^{Venus/+}$ (con) and $Foxa2^{Venus/Venus}$ (ko) endoderm differentiating cells. Dashed regions indicate Foxa2 binding sites.

L) Genome browser view of example transient, stable and late Foxa2 binding sites. The following tracks are displayed: Foxa2 ChIP-seq in d3F and d5FS cells; ATAC-seq in d0, d3F, d5FS cells; H3K27ac ChIP-seq in d0, d3F, d5FS cells; H3K4me1 ChIP-seq in d0, d3F, d5FS cells. Dashed regions indicate Foxa2 binding sites.

**Supplement related to Figure 5**

A) Genome browser view of example stable and late Foxa2 binding sites. The following tracks are displayed: Foxa2 ChIP-seq and Gata4 ChIP-seq in d3F and d5FS cells; ATAC-seq, H3K27ac ChIP-seq and H3K4me1 ChIP-seq in d0, d3F, d5FS cells. Dashed regions indicate Foxa2 binding sites.

B) Scatter plots showing normalized ATAC-seq signals in d0 vs. d5FS cells at Foxa2/Gata4 co-bound sites. Significant chromatin accessibility changes (padj < 0.05, log2 fold change > 1) are coloured in red.

**Supplement related to Figure 6**

A-C) Genome browser view showing Foxa2 ChIP-seq tracks in endoderm and beta cells, Nanog ChIP-seq in day0 cells and indicated chromatin modifications in d0 cells. Dashed areas show a Nanog binding site in day0 cells (A), a Foxa2 binding site in endoderm (B), a Foxa2 binding site in beta-cells (C). Green and red indicate active and repressive chromatin marks, respectively.

D) Read-density heat map showing the normalized coverage of the following features at Foxa2 transient binding sites (from top): Foxa2 ChIP-seq in d3F, d5FS and beta cells; H3K4me1, H3K27ac, H3K4me3 ChIP-seq and 5hmC meDIP-seq in d0 cells.

E) Read-density heat map showing the normalized coverage of the following features at Foxa2 stable binding sites (from top): Foxa2 ChIP-seq in d3F, d5FS and beta cells; H3K4me1, H3K27ac, H3K4me3 ChIP-seq and 5hmC meDIP-seq in d0 cells .

F) Read-density heat map showing the normalized coverage of the following features at Foxa2 late binding sites (from top): Foxa2 ChIP-seq in d3F, d5FS and beta cells; H3K4me1, H3K27ac, H3K4me3 ChIP-seq and 5hmC meDIP-seq in d0 cells .

G) Read-density heat map showing the normalized coverage of the following features at Foxa2 beta cell binding sites (from top): Foxa2 ChIP-seq in d3F, d5FS and beta cells; H3K4me1, H3K27ac, H3K4me3 ChIP-seq and 5hmC meDIP-seq in d0 cells .

H) Box plot showing the normalized coverage of the indicated chromatin modifications at different categories of Foxa2 binding sites in d0 cells. Transient, stable and late binding sites feature higher levels of active modifications. Wilcoxon ranks-sum test statistics is shown in Table S7.

I) Box plots showing normalized ChIP-seq coverage of active (H3K4me1, H3K27ac, H3K4me3) and repressive (H3K27me3) chromatin marks for endoderm vs. liver FOXA2 binding sites in human ESCs. Endoderm binding sites feature higher levels of active modifications. Wilcoxon ranks-sum test statistics is shown in Table S7.

**Supplement related to Figure 7**

A) Representative FACS plots of $ES^{iFVF}$ cells in non-induced (no doxycycline) and induced (plus doxycycline) conditions.

B) Average expression levels of endoderm marker genes detected by RNA-seq in d0 cells, d5FS cells and d2-FVFp cells. TPM: Transcripts Per Kilobase Million. Error bars depict standard deviation (n=2).

C) Box plot showing normalized H3K4me1 coverage in d0 and d2-FVFp cells at Foxa2 binding sites in d2-FVFp cells. Wilcoxon ranks-sum test statistics is shown in Table S7.

D) Box plot showing normalized H3K27ac coverage in d0 and d2-FVFp cells at Foxa2 binding sites in d2-FVFp cells. Wilcoxon ranks-sum test statistics is shown in Table S7.

E) Box plot showing normalized ATAC-seq coverage of d2-FVFp Foxa2 binding sites in do ESCs (ATAC UP – increased chromatin accessibility in d2-FVFp cells; ATAC NC – no change in chromatin accessibility in d2-FVFp cells). Wilcoxon ranks-sum test statistics is shown in Table S7.

F) Jun-AP1 motif density at d2-FVFp Foxa2 binding sites with (ATAC UP) or without (ATAC NC) changes in chromatin accessibility.

G) Representative FACS plots of $ES^{iFVF\text{-}Gata}$ cells in non-induced (no doxycycline) and induced (plus doxycycline) conditions.

### Supplement related to Figure 1

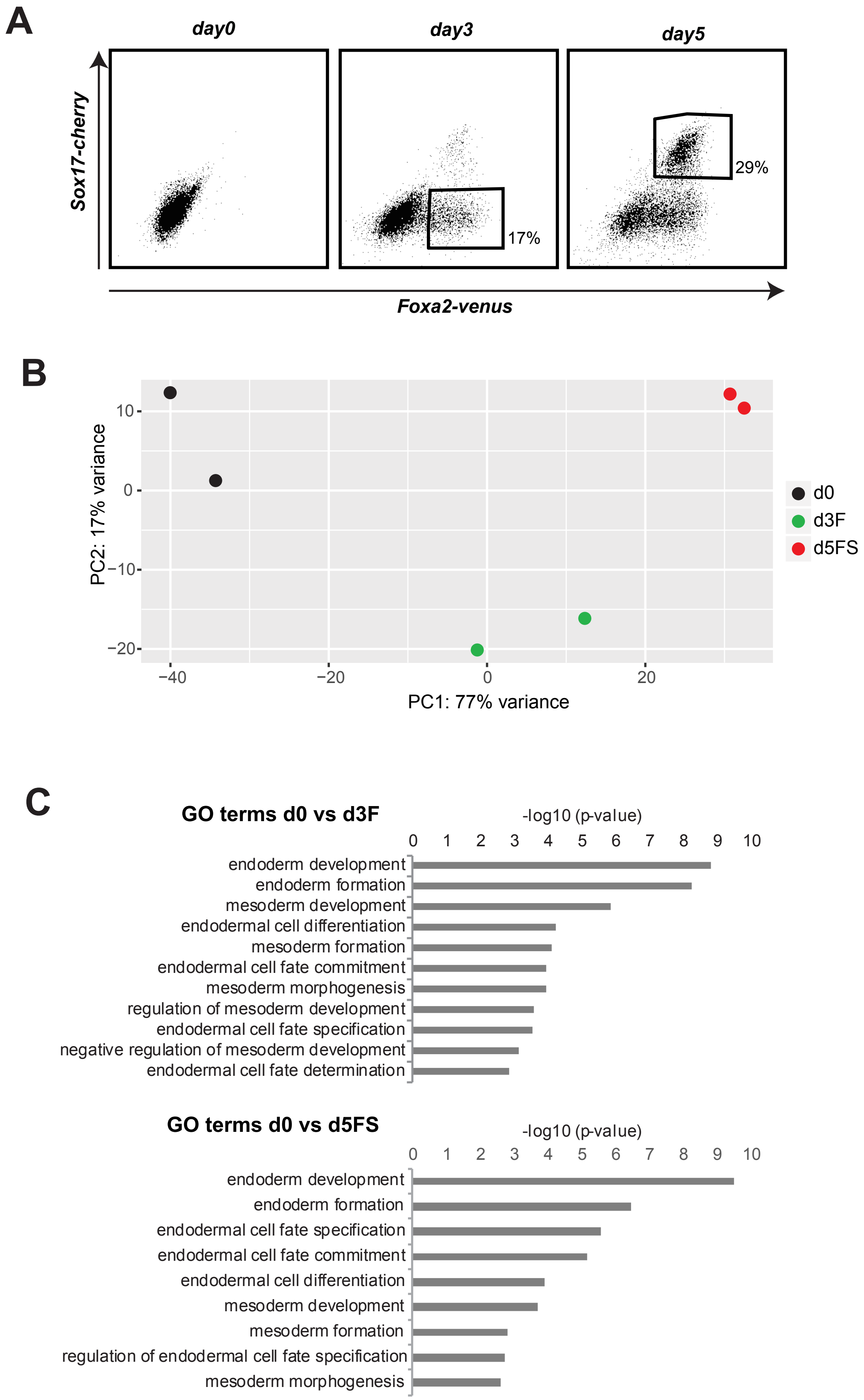

### Supplement related to Figure2

**A**

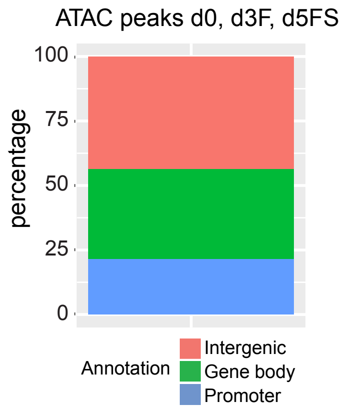


**B**

Gene expression and ATAC changes

| transition | % genes UP | % genes DOWN |
|---|---|---|
| d0-d3F | 19 | 46 |
| d3F-d5FS | 21 | 32 |
| d0-d5FS | 45 | 78 |

**C**

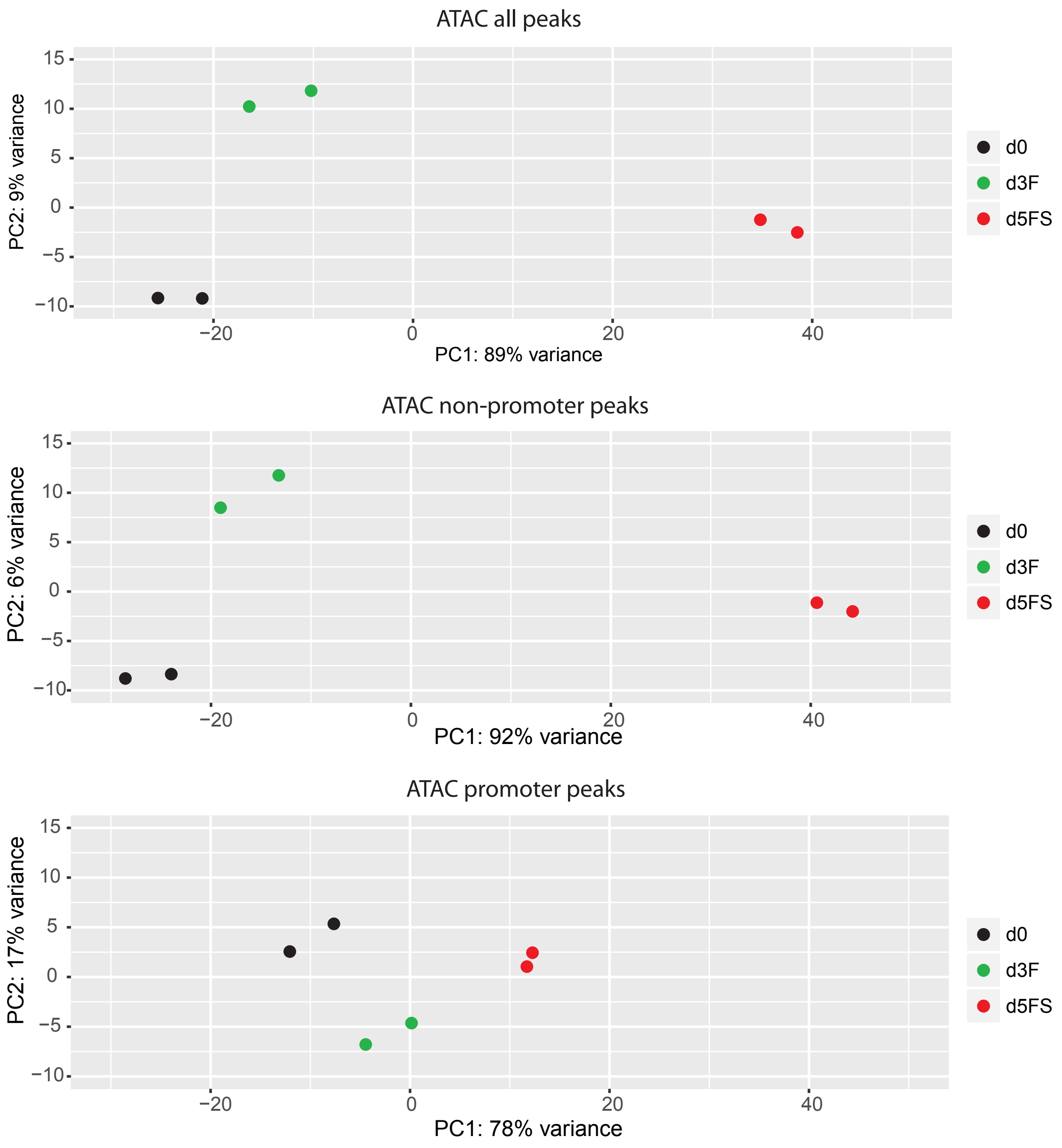

### Supplement related to Figure2

**D**

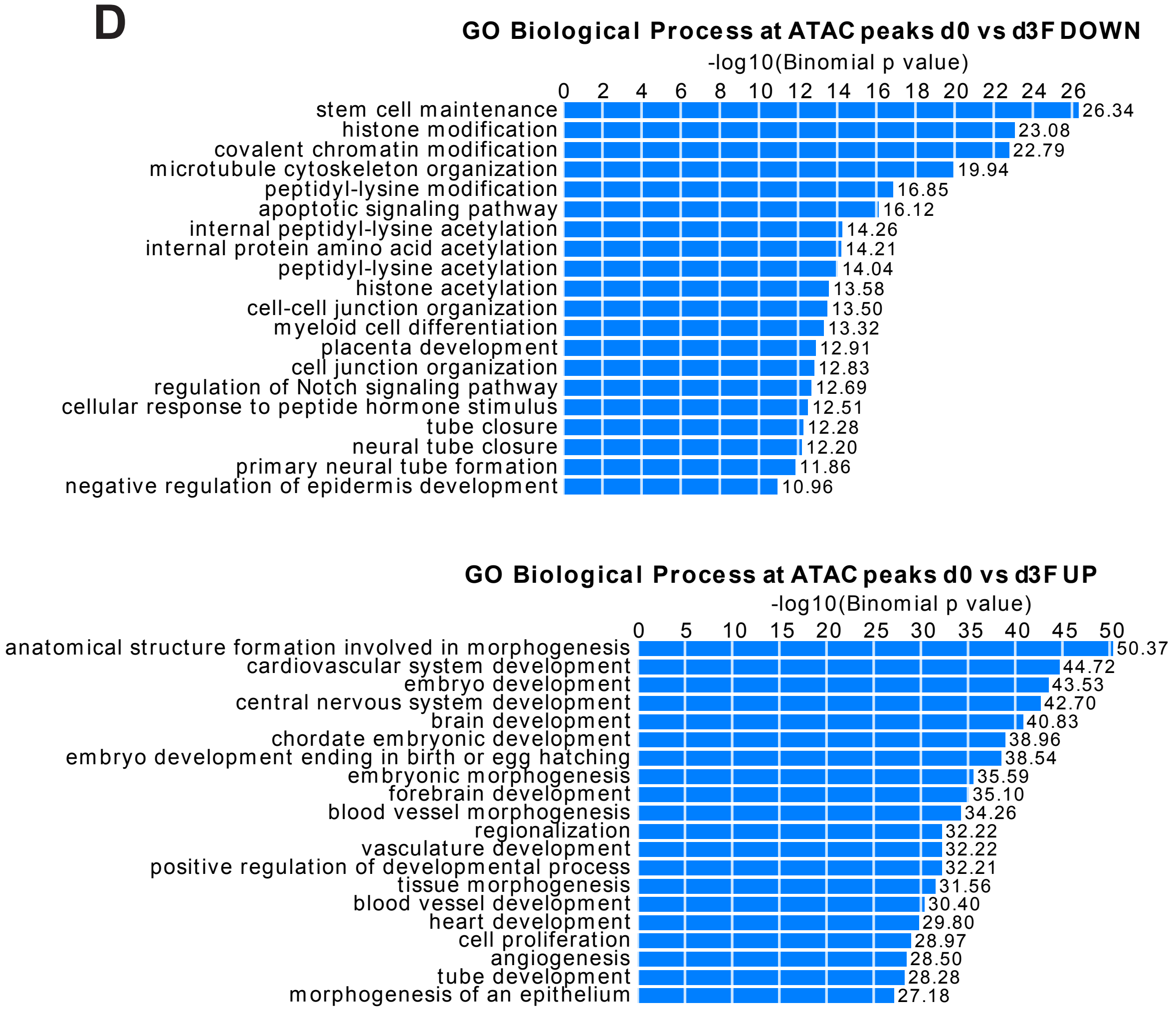


**E**

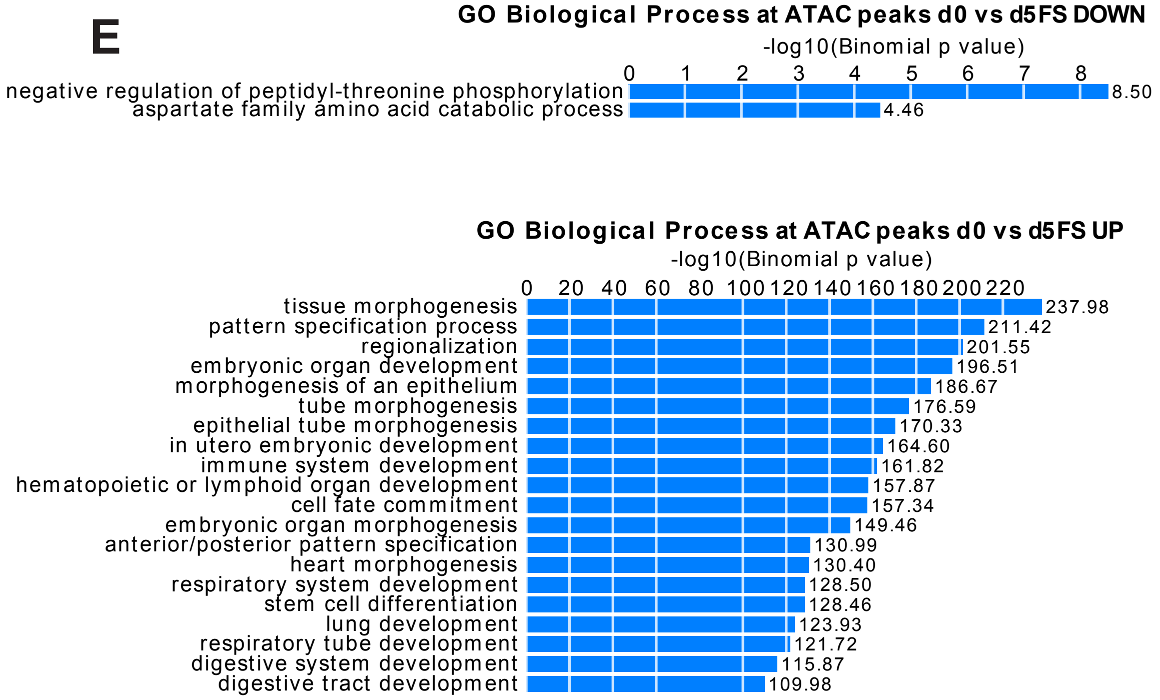

### Supplement related to Figure2

**F**

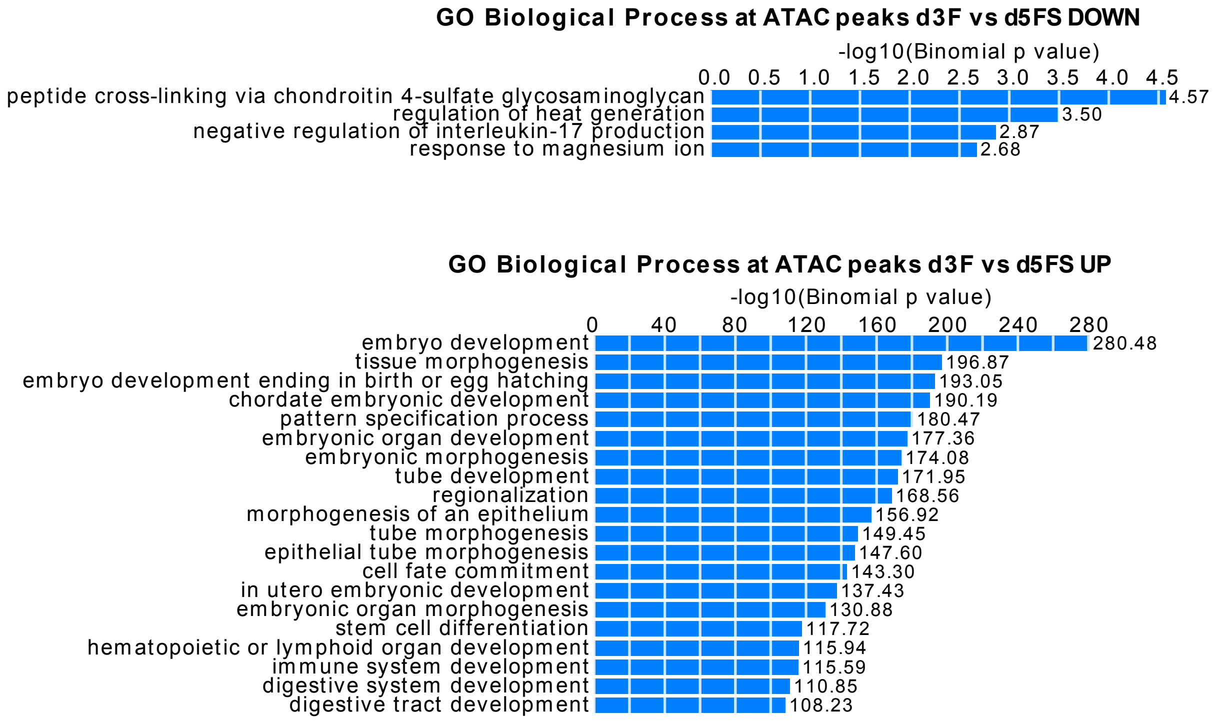

### Supplement related to Figure 3

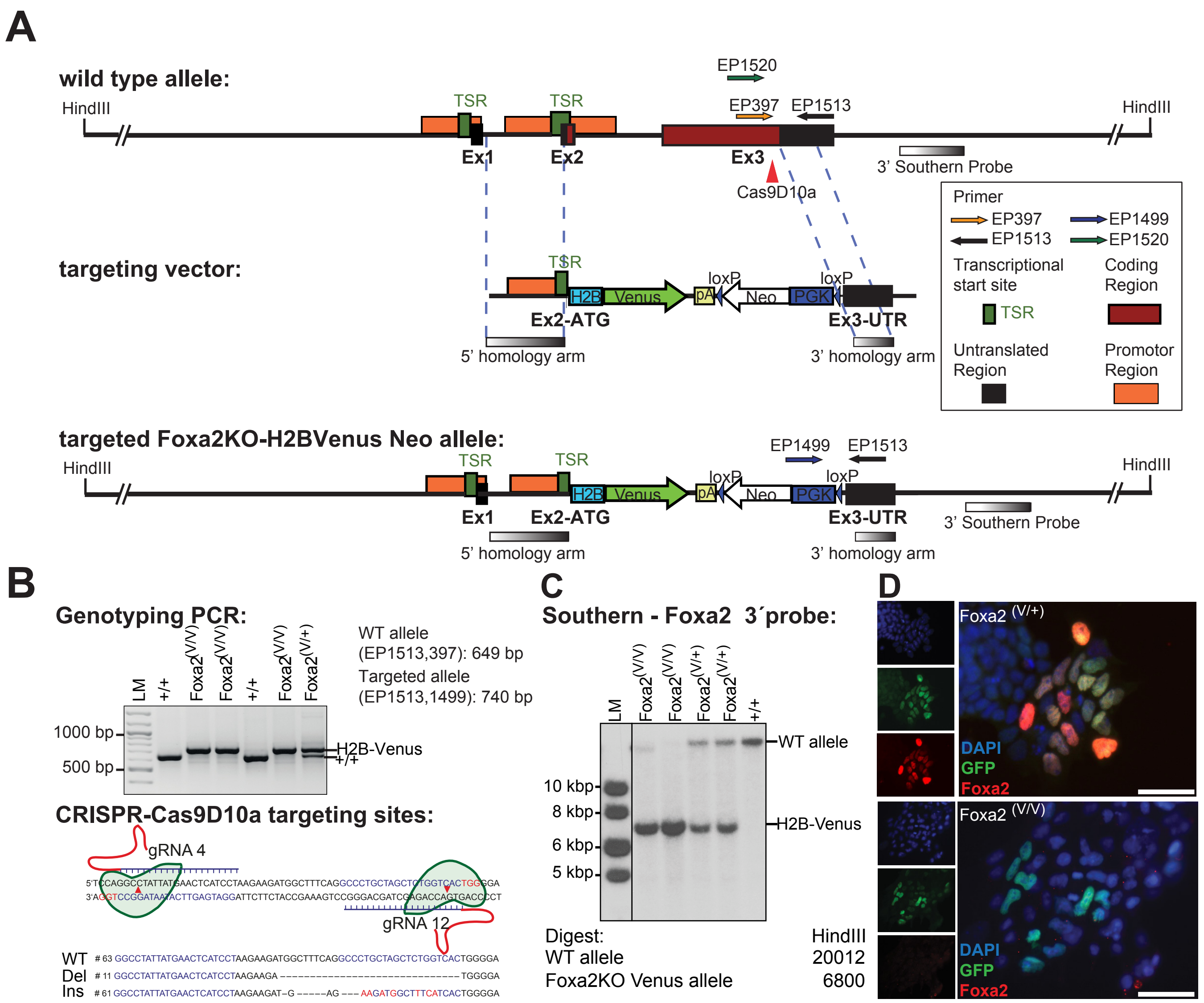

#### Supplement related to Figure3

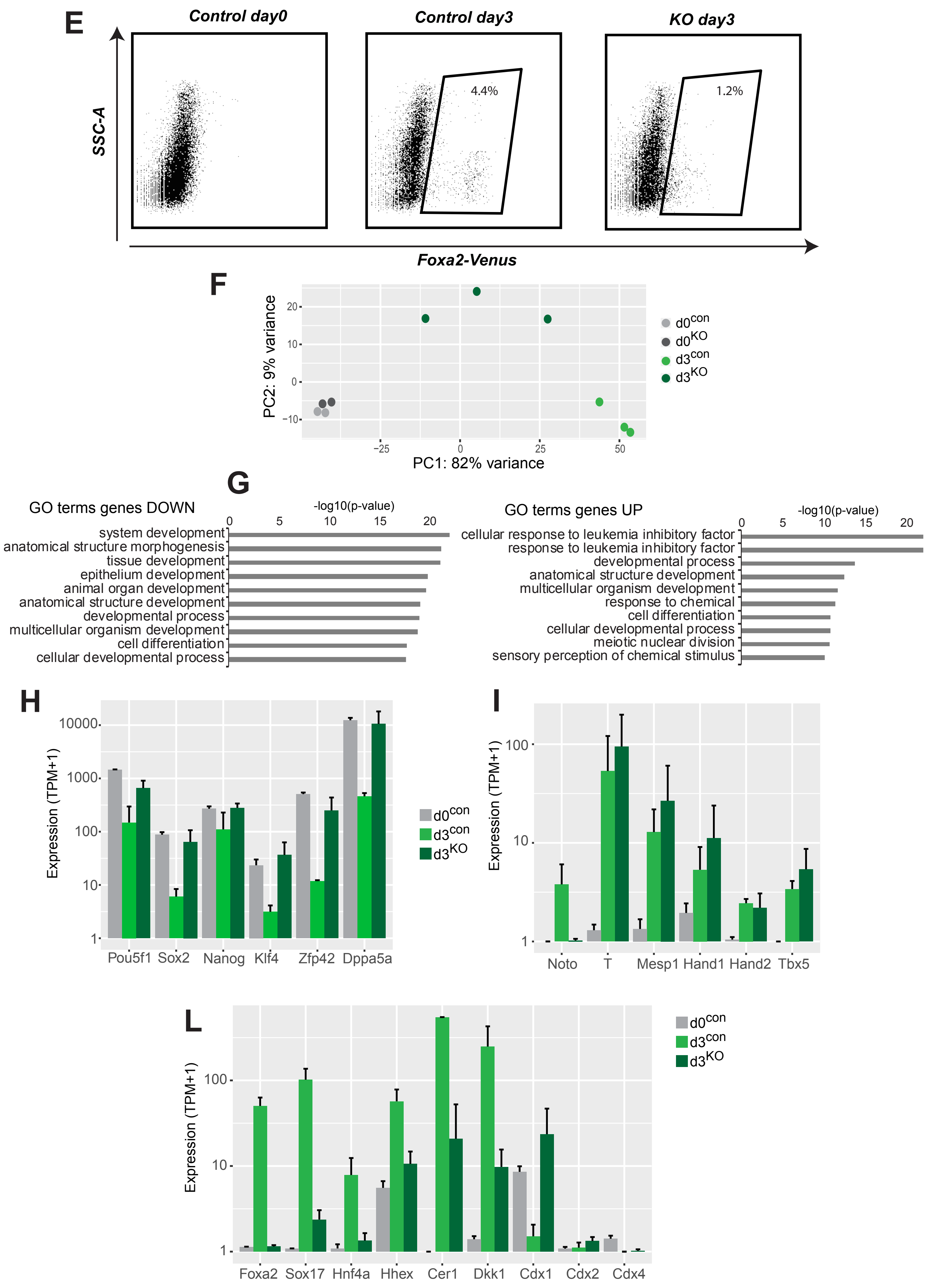

### Supplement related to Figure 4

**A**

##### Top motifs at Transient Foxa2 peaks

| Rank | Homer Known Motif | Name | p-val | % of targets |
|---|---|---|---|---|
| 1 | 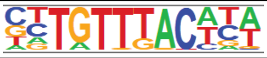 | Foxa2 | 1e-1655 | 70.86 |

##### Top motifs at Stable Foxa2 peaks

| Rank | Homer Known Motif | Name | p-val | % of targets |
|---|---|---|---|---|
| 1 | 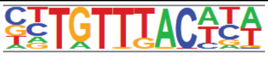 | Foxa2 | 1e-3037 | 82.70 |

##### Top motifs at Late Foxa2 peaks

| Rank | Homer Known Motif | Name | p-val | % of targets |
|---|---|---|---|---|
| 1 | 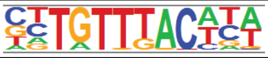 | Foxa2 | 1e-1996 | 74.87 |

**B**

##### GO Biological Process at Transient Foxa2 peaks

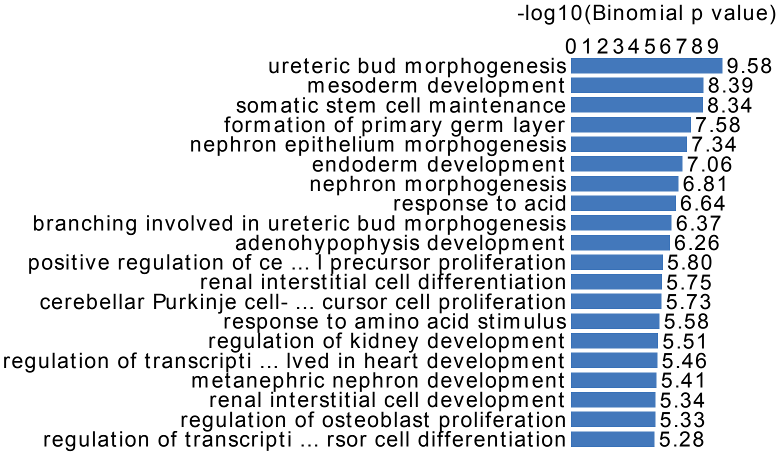


##### GO Biological Process at Stable Foxa2 peaks

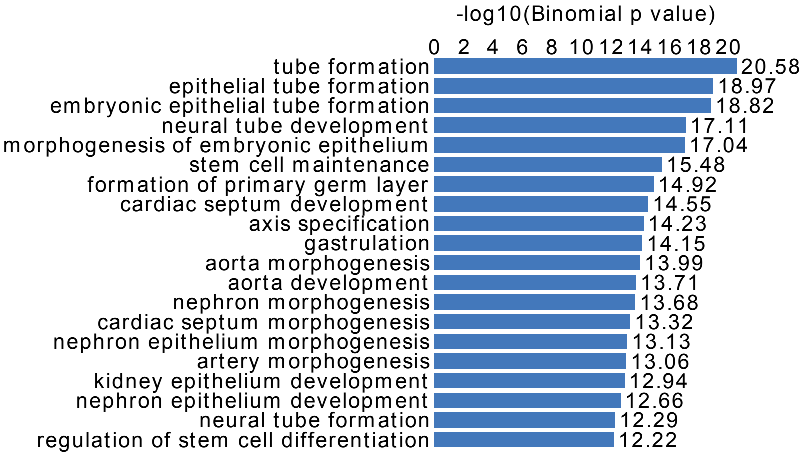


##### GO Biological Process at Late Foxa2 Peaks

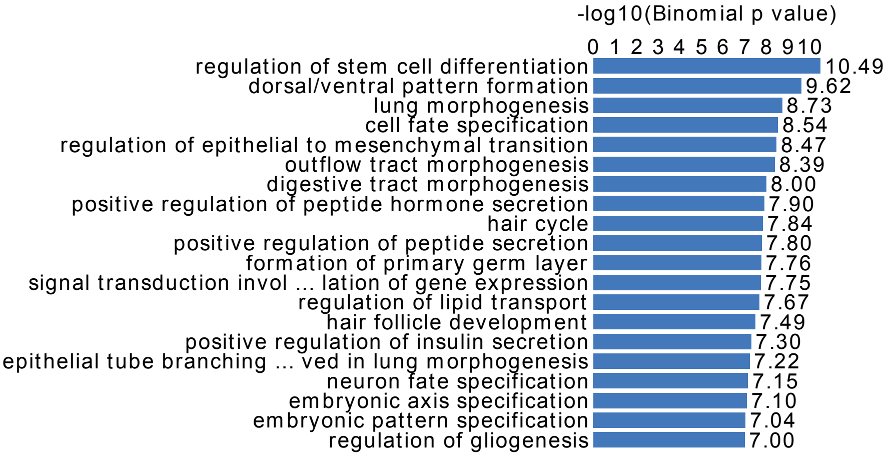

#### Supplement related to Figure 4

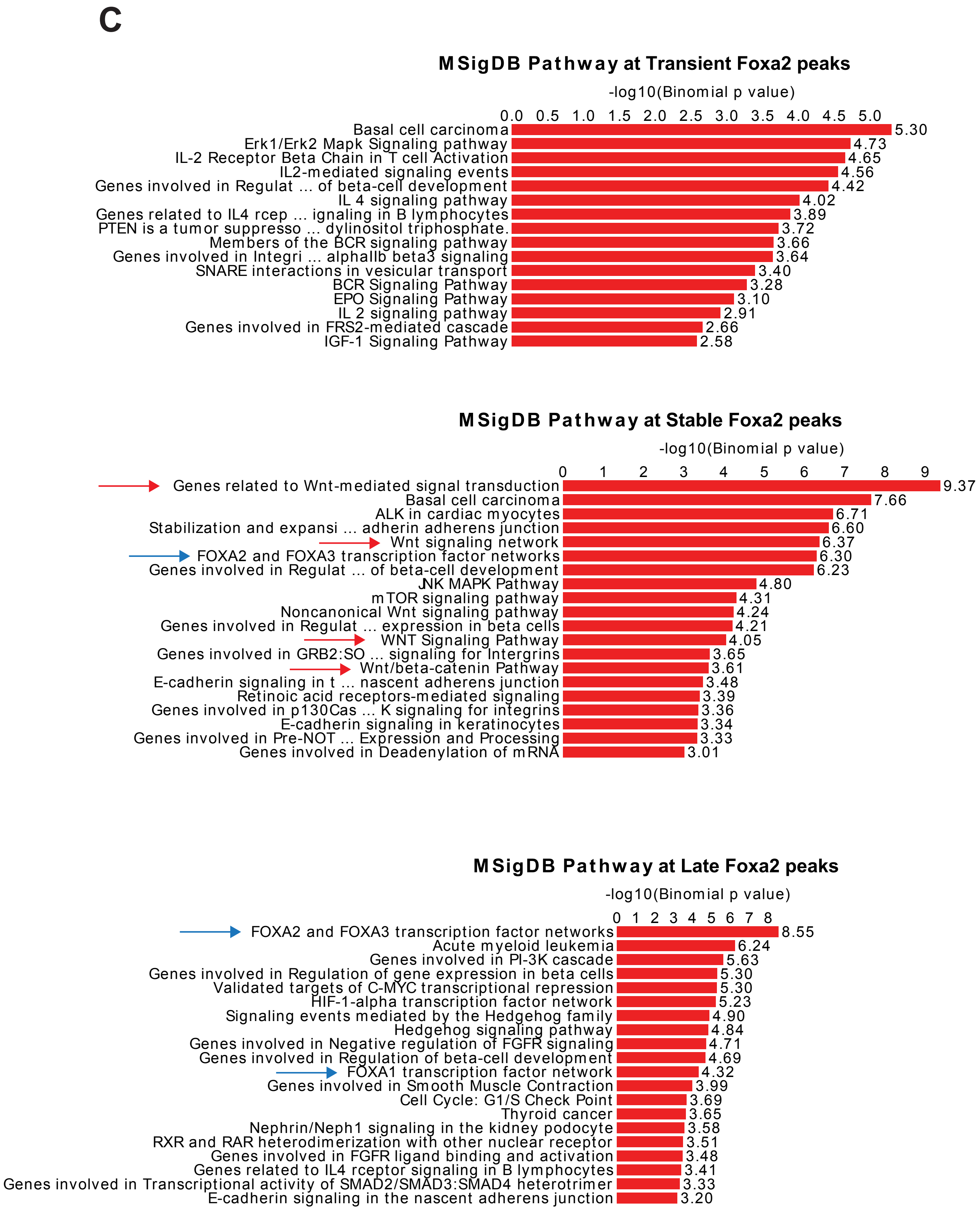

### Supplement related to Figure 4

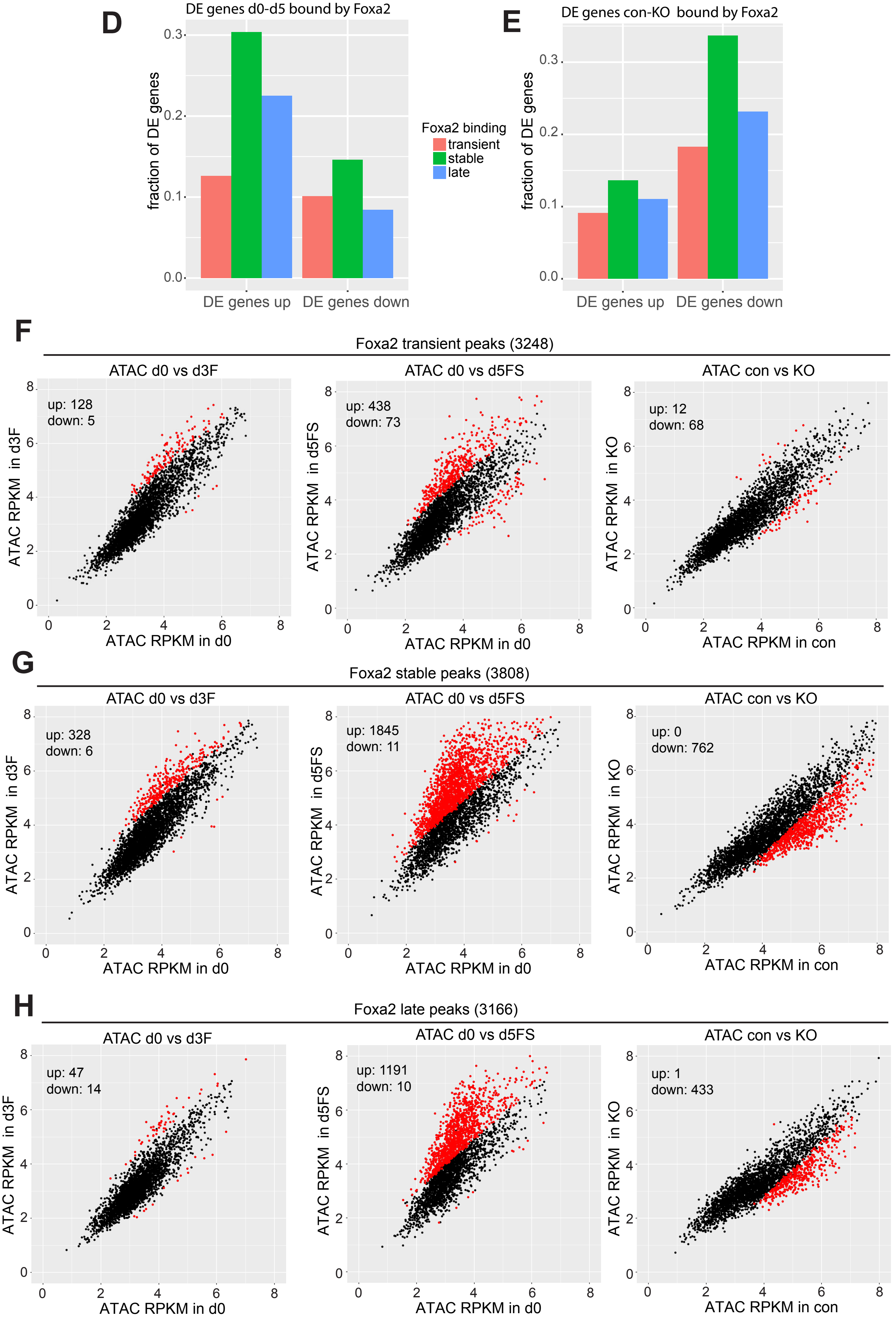

### Supplement related to Figure 4

**I**

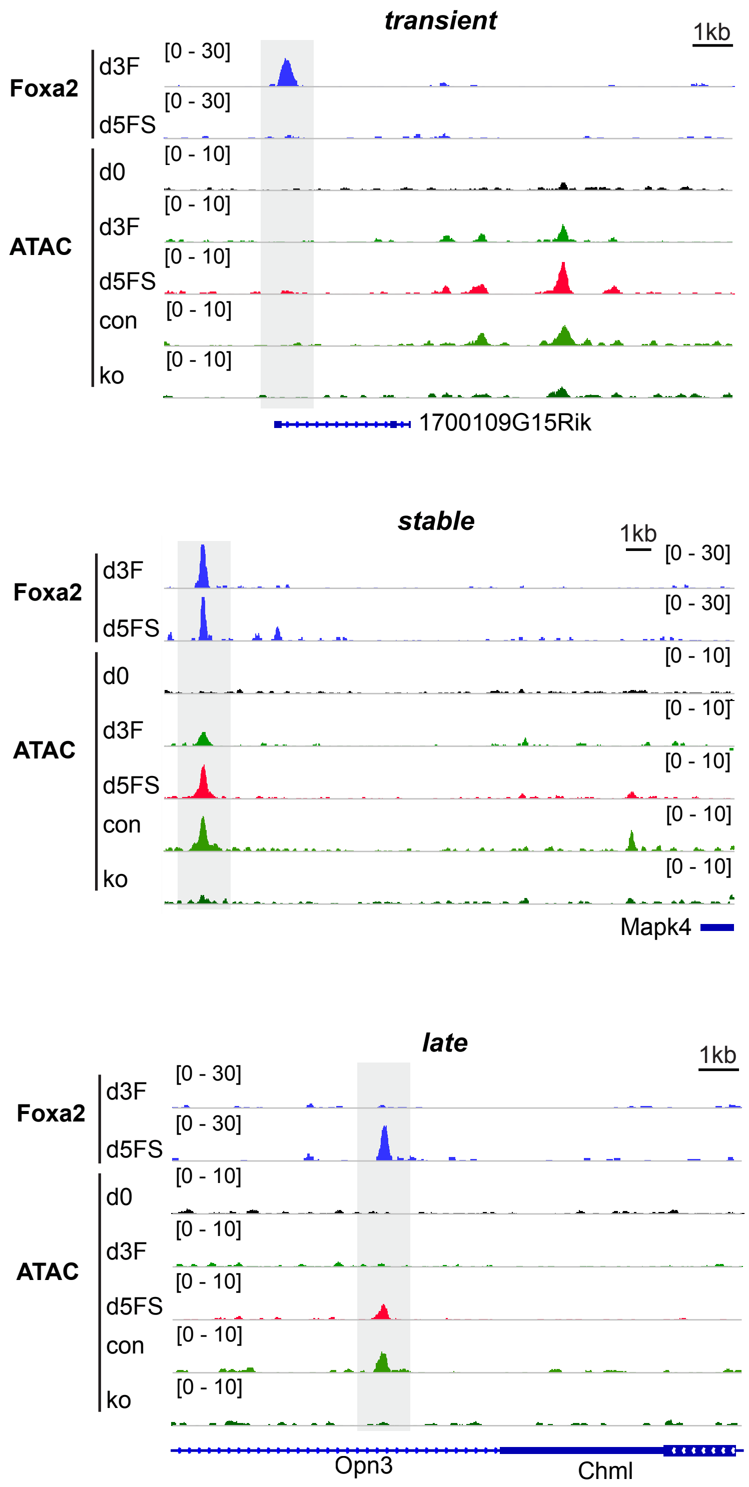

### Supplement related to Figure 4

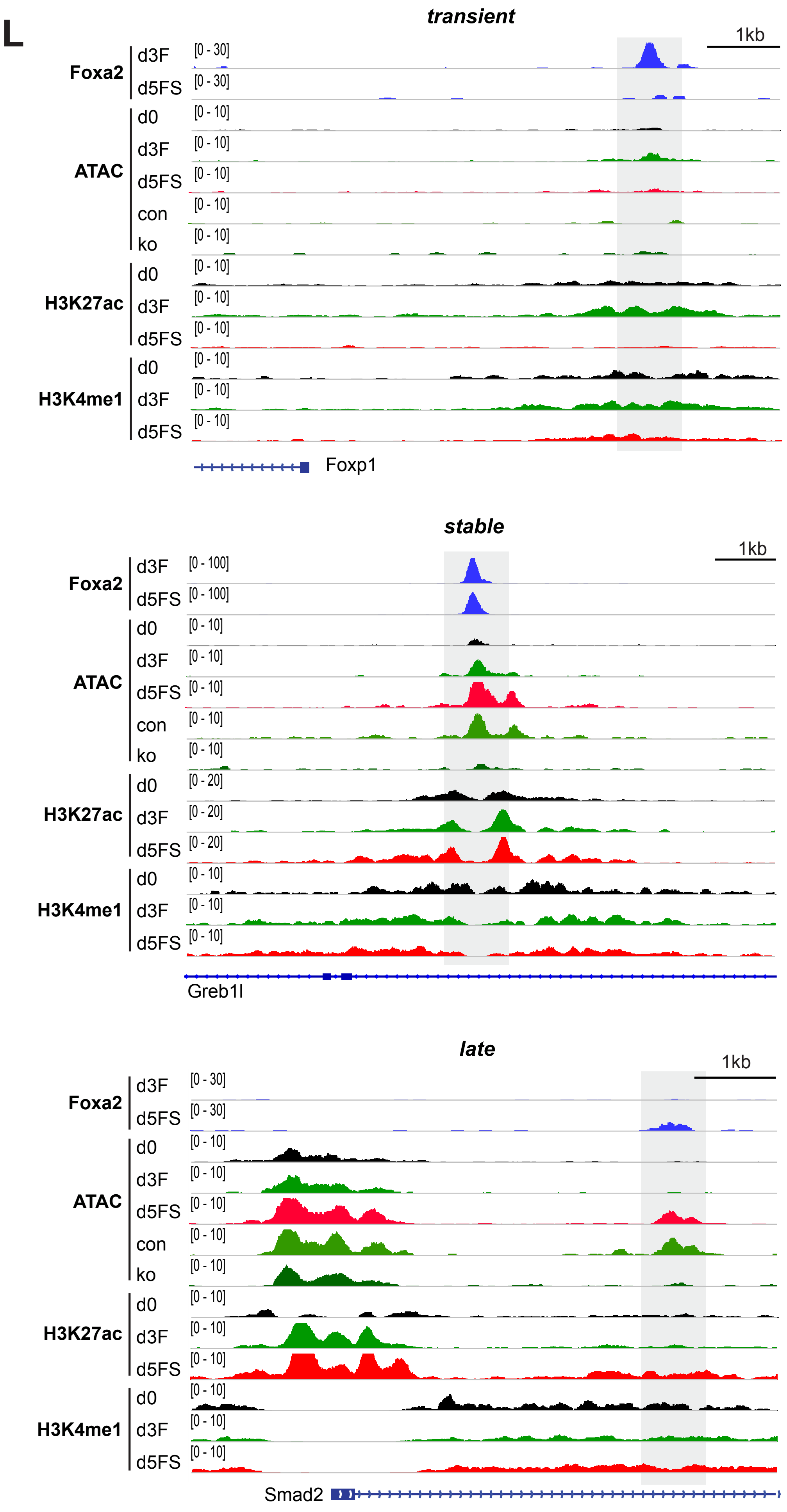

### Supplement related to Figure 5

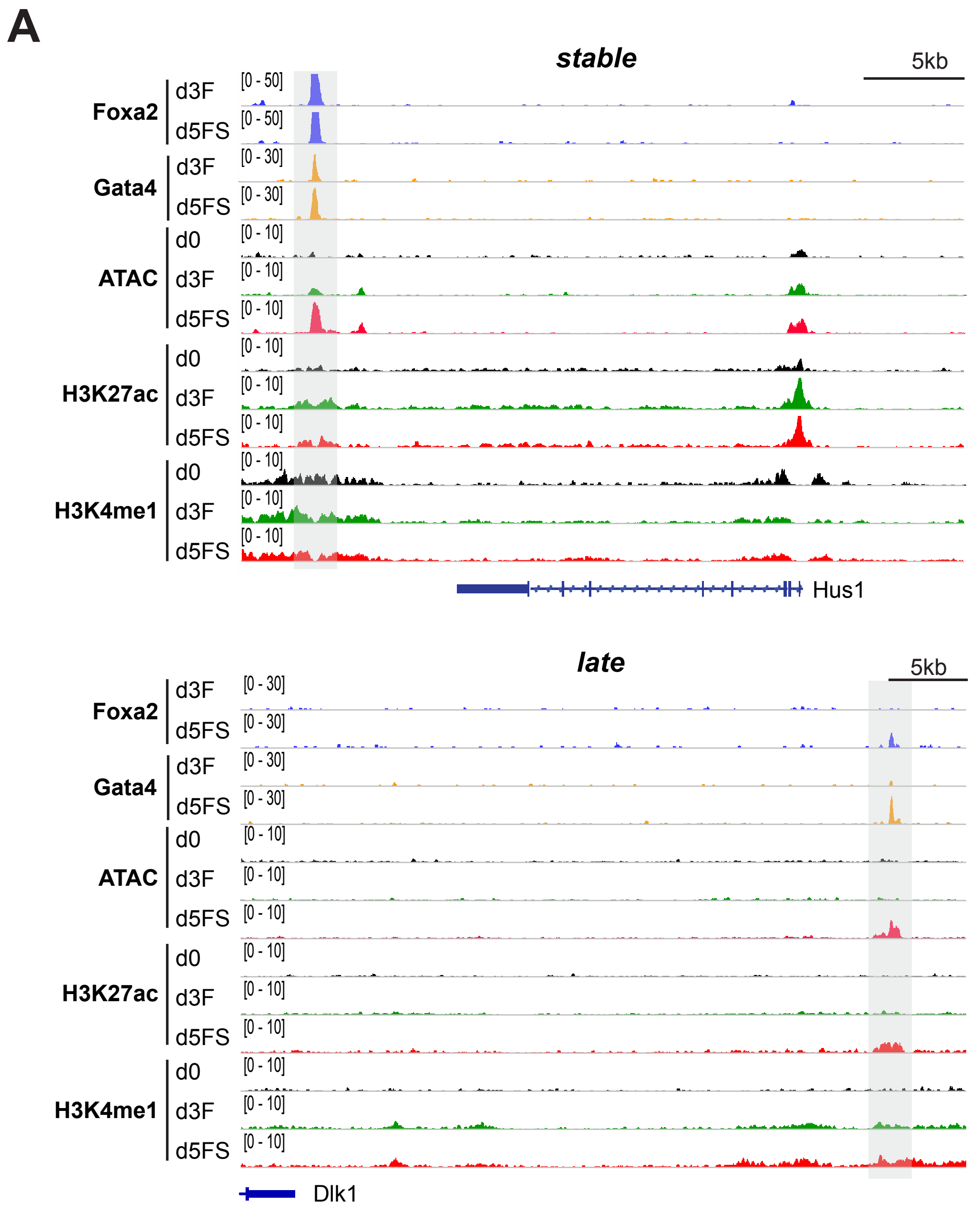


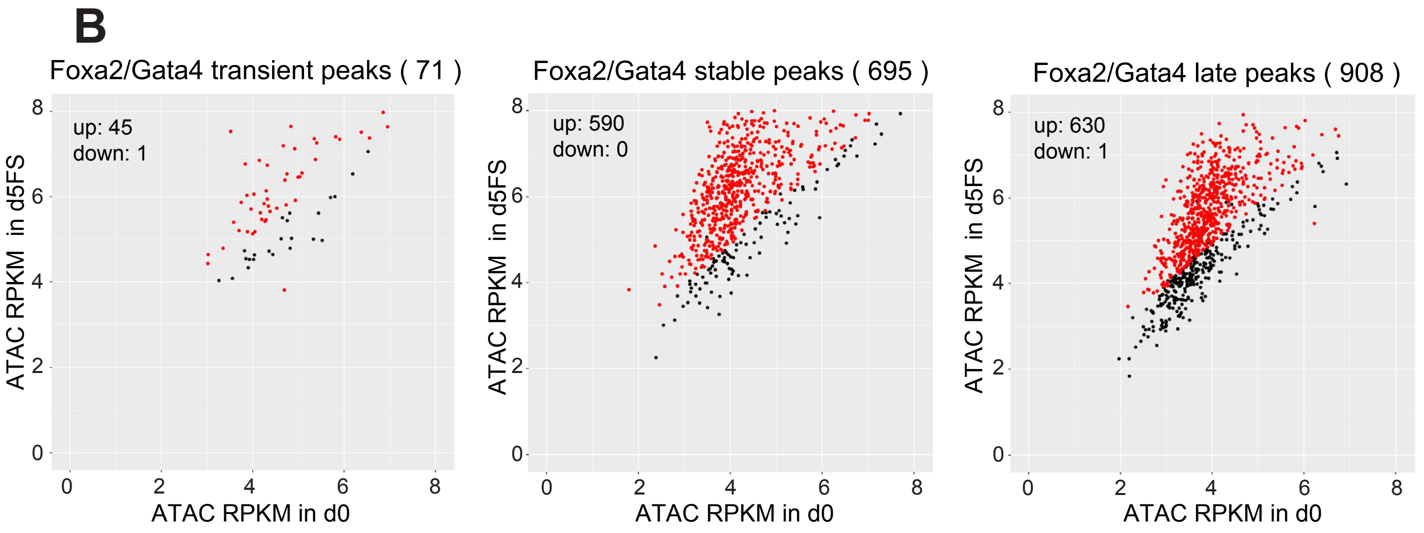

#### Supplement related to Figure 6

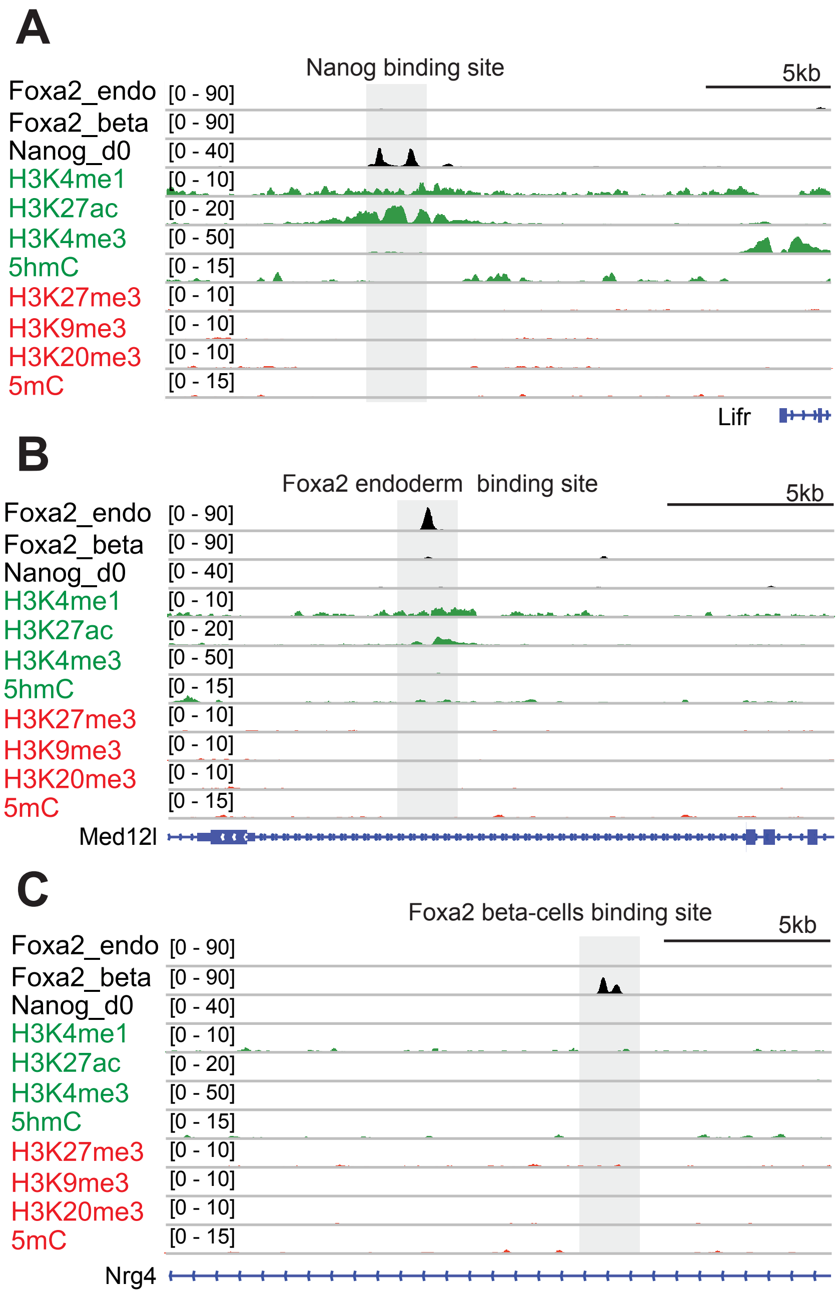

### Supplement related to Figure 6

D

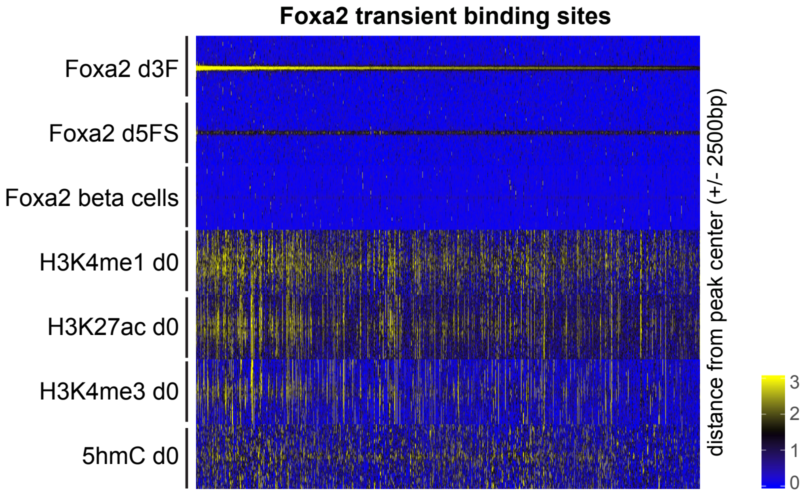


E

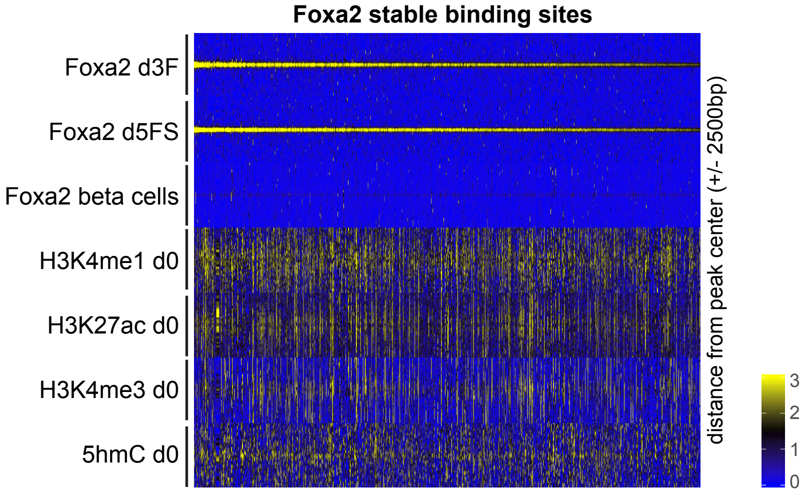


F

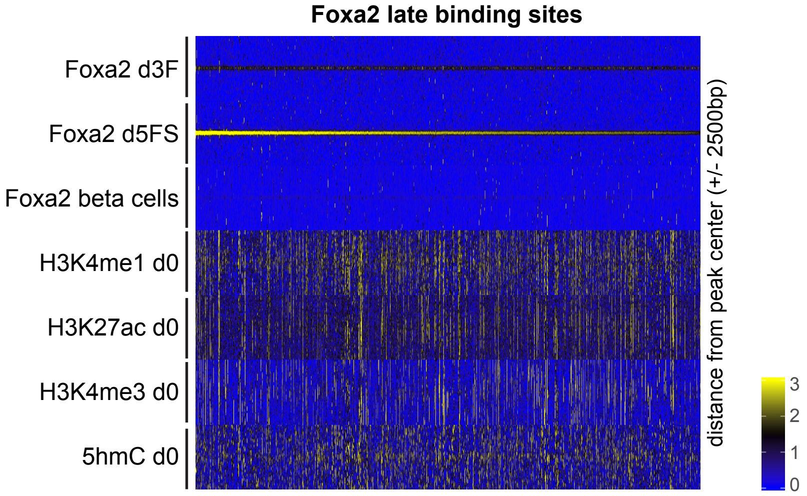


G

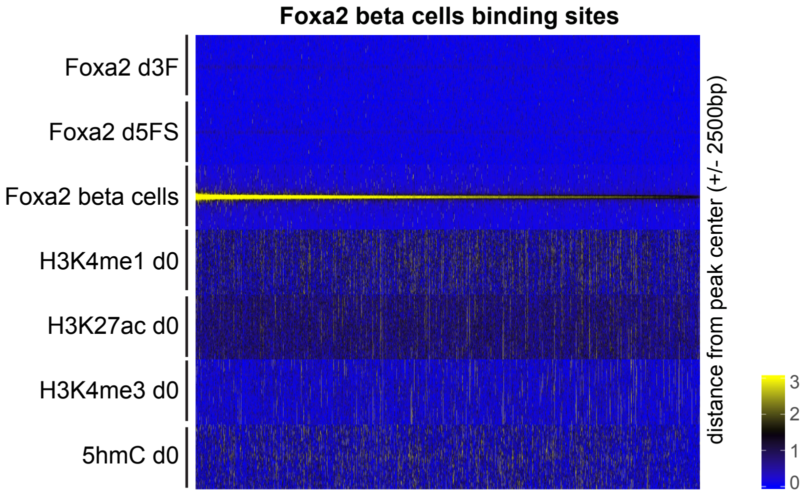

#### Supplement related to Figure 6

**H**

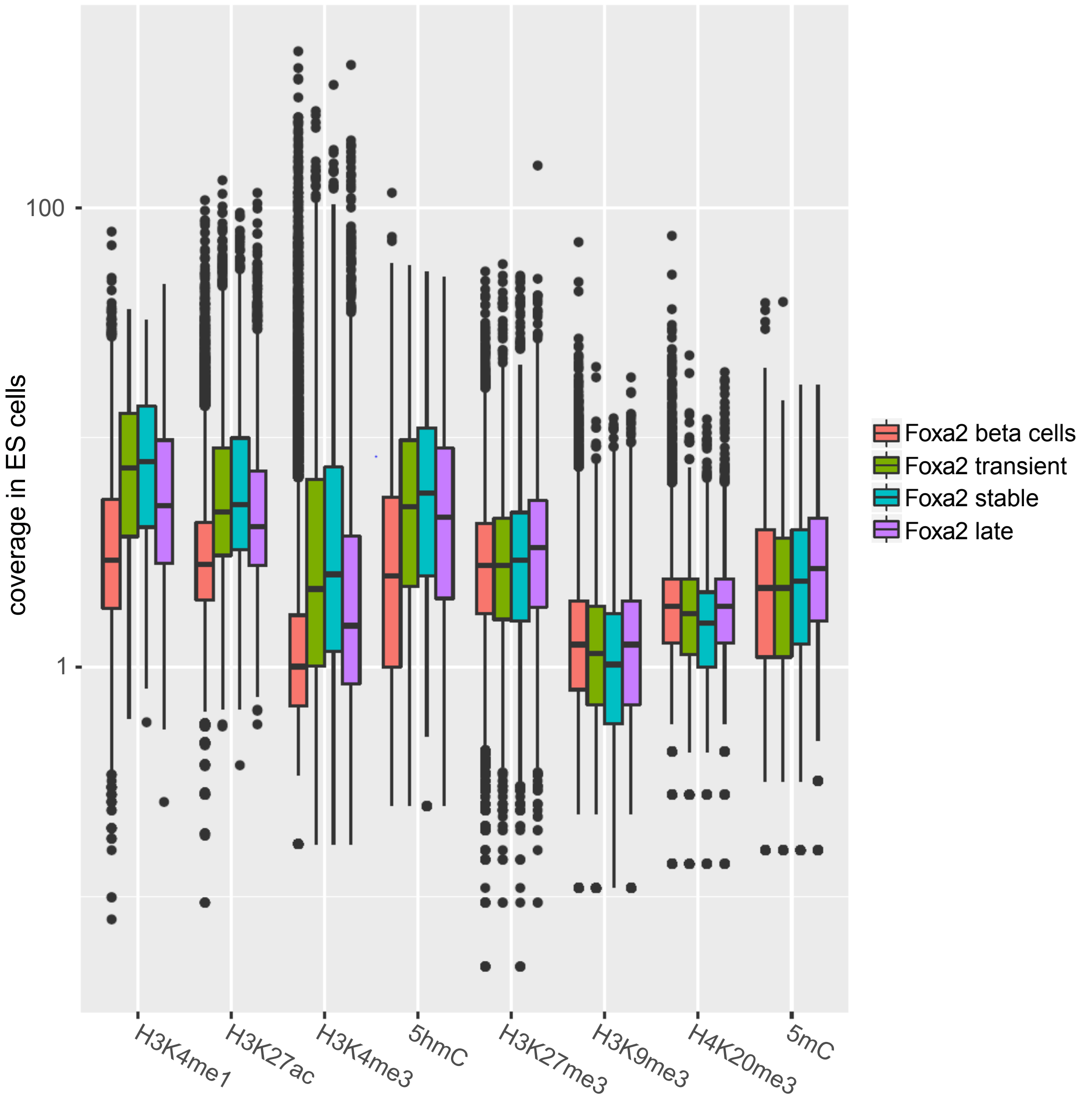


**I**

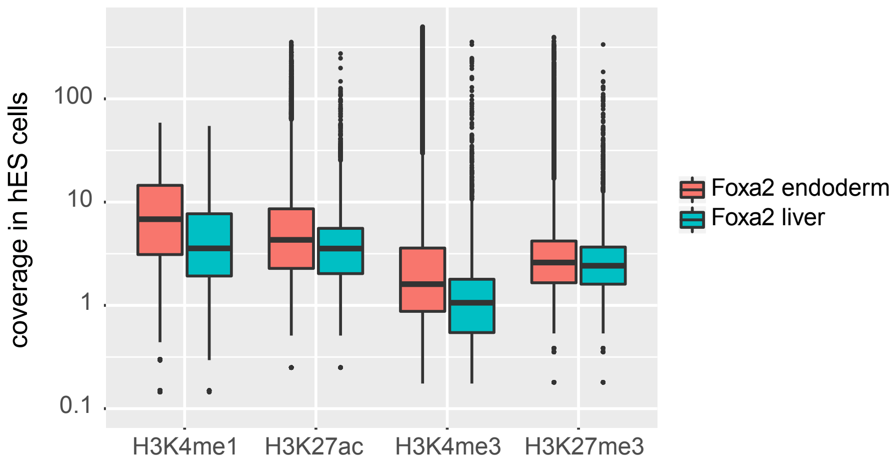

### Supplement related to Figure7

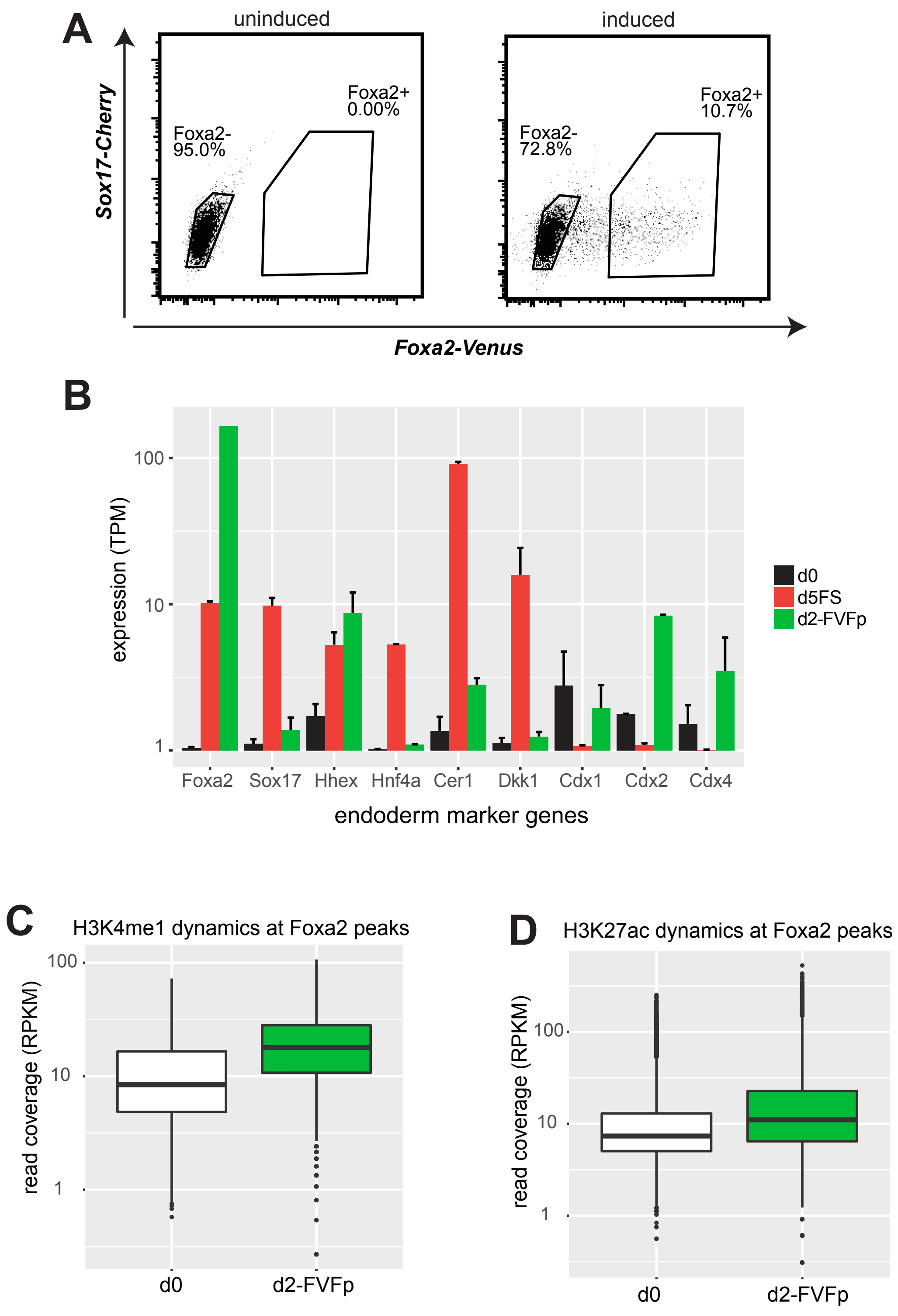

### Supplement related to Figure7

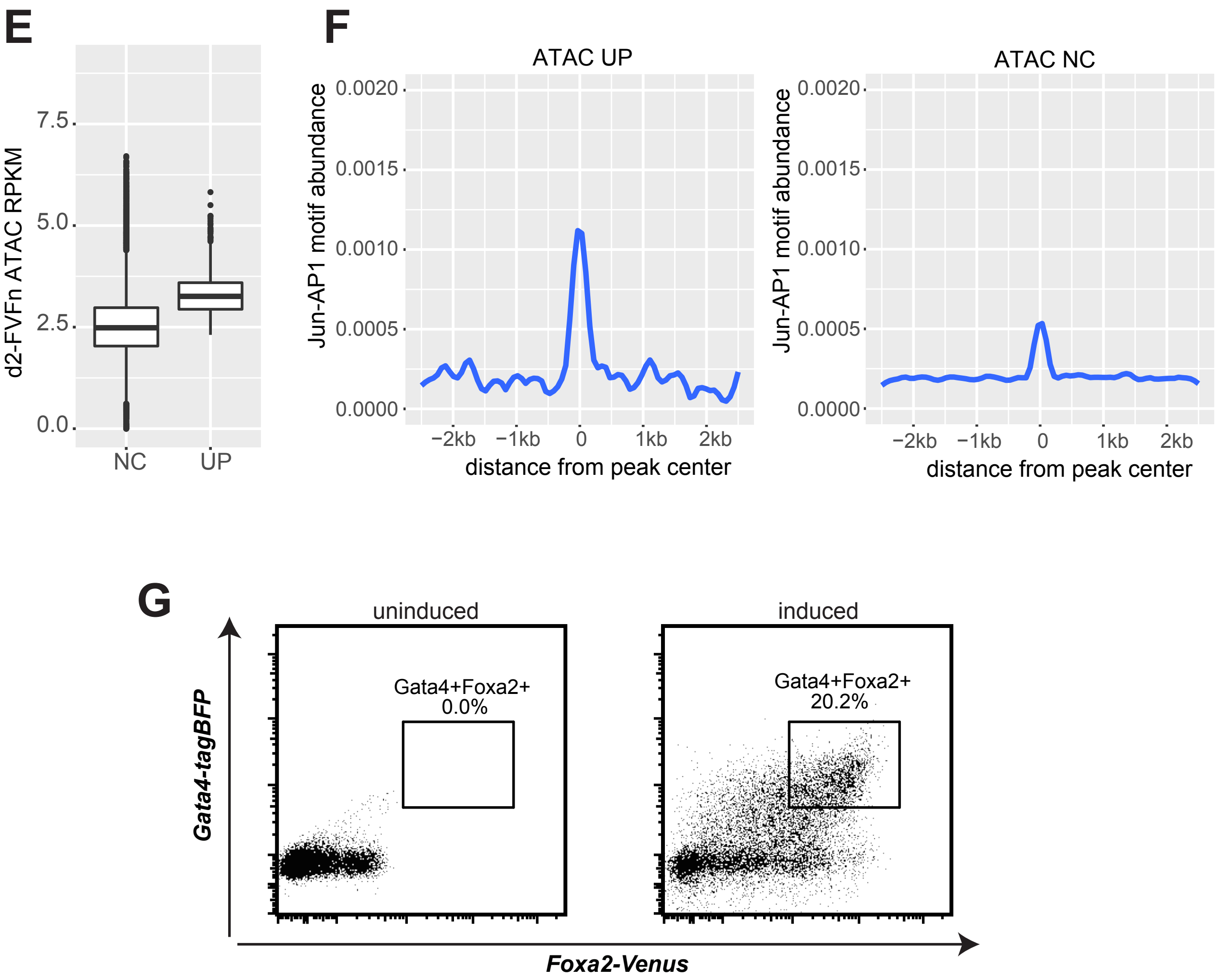
